# Environmental DNA substrates capture complementary dimensions of insect biodiversity

**DOI:** 10.64898/2026.09.13.750568

**Authors:** Beilun Zhao, John Sundh, Tobias Andermann

**Affiliations:** Key Laboratory of Marsh Wetland Ecosystem Conservation and Restoration, National Forestry and Grassland Administration, Key Laboratory of Wetland Ecology, Northeast Institute of Geography and Agroecology, Chinese Academy of Sciences, 130102 Changchun, China; National Field Observation and Research Station (Heilongjiang) for Xingkai Lake Wetland Ecosystem, Northeast Institute of Geography and Agroecology, Chinese Academy of Sciences, 130102 Changchun, China; Department of Organismal Biology, Uppsala University, Uppsala, Sweden; Science for Life Laboratory, Uppsala University, Uppsala, Sweden; Dept of Biochemistry and Biophysics, National Bioinformatics Infrastructure Sweden, Science for Life Laboratory, Stockholm University, Stockholm, Sweden

**Keywords:** environmental DNA, metabarcoding, insect monitoring, biodiversity, substrate complementarity

## Abstract

Environmental DNA (eDNA) metabarcoding has become a powerful approach for biodiversity assessment, yet whether different environmental substrates provide equivalent or complementary representations of terrestrial biodiversity remains unresolved. Here we addressed this question by comparing insect assemblages recovered from Malaise traps and six environmental substrates (water, sediment, soil, spiderwebs, deadwood, and anthills) across 45 forest locations using a standardized COI metabarcoding workflow. Substrate identity consistently shaped insect detection, diversity, taxonomic representation, and community composition, with each substrate recovering distinct components of insect biodiversity rather than a common biodiversity signal. Malaise traps recovered the greatest overall richness, while water and spiderwebs contributed the largest proportion of additional taxa, demonstrating that no single substrate represented the full spectrum of insect biodiversity. These findings show that environmental substrates capture complementary representations of biodiversity through distinct ecological processes of eDNA deposition, transport, accumulation, and persistence. Different substrates should therefore be interpreted as complementary sources of biodiversity observations instead of interchangeable sampling media, and their integration can support more comprehensive ecological inference. Our study identifies substrate complementarity as a key consideration for interpreting terrestrial eDNA observations and for designing future biodiversity surveys.

## 1 Introduction

Insects constitute the most diverse group of terrestrial animals and play essential roles in ecosystem functioning through pollination, decomposition, nutrient cycling, herbivory, and food-web interactions (May, 1986; Scudder, 2017; Tihelka et al., 2021). Growing evidence, however, indicates widespread declines in insect abundance, biomass, and diversity across many regions of the world (Johnson & Haynes, 2023; Outhwaite et al., 2022). Detecting and understanding these changes requires biodiversity monitoring methods that are scalable, reproducible, and taxonomically comprehensive. Conventional insect surveys, including Malaise traps (MT), pitfall traps, light traps, and direct observations, have long provided the foundation for ecological monitoring but remain labor-intensive, taxonomically demanding, and difficult to standardize across broad spatial and temporal scales (Kirse et al., 2021; Marquina et al., 2019).

Environmental DNA (eDNA) metabarcoding has emerged as a powerful alternative for biodiversity assessment by enabling species detection from DNA traces released into the environment (Bohmann & Lynggaard, 2023; Thomsen & Willerslev, 2015). Originally developed for aquatic ecosystems, eDNA is now increasingly applied to terrestrial biodiversity monitoring, including insects (Gregorič et al., 2022; Guthrie et al., 2023; Kirse et al., 2021). Unlike conventional trapping, terrestrial eDNA can be recovered from a wide variety of environmental substrates, including water, soil, sediments, spiderwebs, deadwood, and other natural matrices that accumulate biological material through different physical and ecological processes over time (Ryan et al., 2022; van der Heyde et al., 2020; Zhao & Andermann, 2026). These approaches expand opportunities for standardized biodiversity surveys while increasing the likelihood of detecting taxa that are difficult to capture using conventional sampling alone.

Despite rapid methodological development, the extent to which different environmental substrates capture overlapping or complementary biodiversity information remains poorly understood. Previous studies have demonstrated that individual substrates can successfully recover diverse insect assemblages, including freshwater communities from water and sediment samples (Takenaka et al., 2024), terrestrial arthropods from soil (Lunghi et al., 2022), and airborne biodiversity signals from spiderwebs (Gregorič et al., 2022). However, most comparisons have been limited to one or two environmental matrices or a small number of sampling locations (Banerjee et al., 2026; Karlsson et al., 2020; Miraldo et al., 2025), making it difficult to determine whether different substrates recover largely redundant biodiversity signals or consistently capture distinct dimensions of insect communities. Resolving this question is essential because it determines how eDNA observations should be interpreted and whether integrating multiple substrates provides a more complete representation of biodiversity than any single substrate.

Environmental substrates differ fundamentally in the processes governing DNA deposition, transport, accumulation, and persistence (Bell et al., 2024; Nagler et al., 2022; van der Heyde et al., 2022). Consequently, they are expected to preserve different ecological information and represent different dimensions of insect biodiversity. Flowing water integrates DNA signals across landscapes through hydrological connectivity (Egeter et al., 2018; Shogren et al., 2018; Yao et al., 2022), whereas soils predominantly retain localized signals shaped by microhabitat conditions (Foucher et al., 2020; Kirse et al., 2021). Spiderwebs accumulate airborne biological material through the interception of small organisms, body fragments, and environmental particles (Gregorič et al., 2022; Newton et al., 2024). Similar differences are expected among other terrestrial substrates. Consequently, each substrate may preferentially retain different components of biodiversity rather than providing a complete representation of local communities (Wu et al., 2026). If so, environmental substrates should be viewed as complementary sources of ecological information, with each capturing different dimensions of insect biodiversity.

To address this knowledge gap, we conducted a large-scale comparison of seven substrate types, including MT samples, water, sediment, soil, spiderwebs, deadwood, and anthills, collected across 45 spruce forest locations in Sweden. Using a standardized metabarcoding workflow, we evaluated substrate performance in terms of insect detection efficiency, alpha diversity, community composition, taxonomic representation, and species overlap. Specifically, we asked: (1) how do different substrates compare in their ability to recover insect diversity; (2) to what extent do substrates recover overlapping versus unique components of insect communities; and (3) how does substrate choice influence ecological interpretations of biodiversity patterns? By integrating MTs and multiple eDNA substrates within a single sampling framework, this study provides one of the most comprehensive assessments of terrestrial insect eDNA substrates to date and offers new insights into how substrate selection shapes biodiversity monitoring outcomes.

## 2 Materials and Methods

### 2.1 Sampling design and sample collection

To systematically compare insect diversity across multiple substrates and locations, we conducted field sampling in southern Sweden from October 2 to November 9, 2023. To reduce ecological variability, all 45 sites were selected in spruce-dominated forest (*Picea abies*) adjacent to water bodies (Table S1; Figure 1), spanning longitudes 12.4°-17.9° and latitudes 56.3°-60.2° (WGS84). To reduce environmental heterogeneity among sites and to facilitate field sampling, we grouped the 45 sites into 15 triplets of nearby locations (e.g., 1A, 1B and 1C).

**Figure 1.**
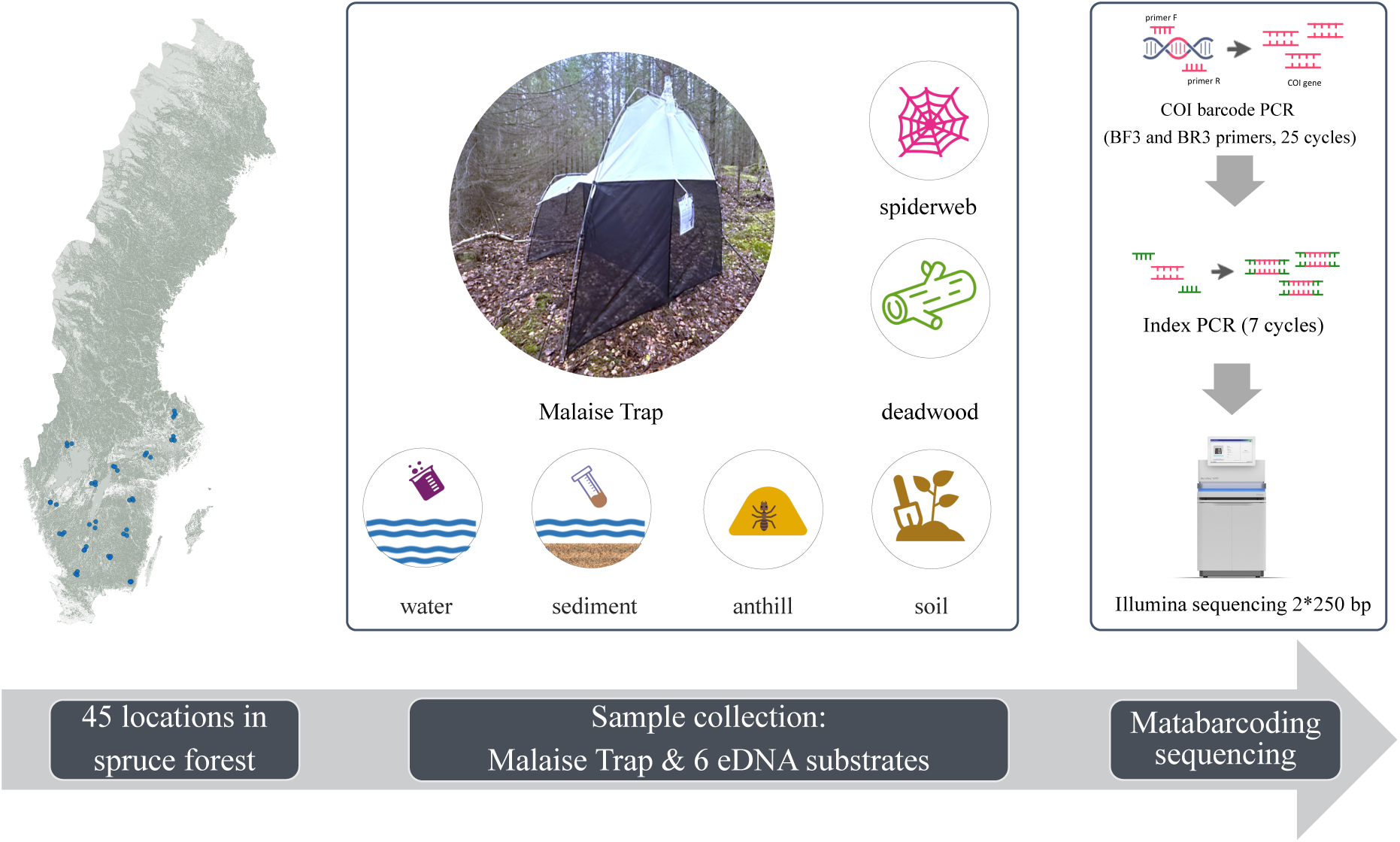
Sample collection and processing. The left panel shows the geographic distribution of the 45 sampling locations across southern Sweden, all situated in spruce forests adjacent to water bodies. At each location, a MT was deployed for one week to collect bulk insect samples. Concurrently, six types of environmental DNA (eDNA) substrates were collected (middle panel). After sample collection, DNA extraction was performed for each sample, followed by a two-step PCR targeting the cytochrome c oxidase subunit I (COI) gene region and adding Illumina sequencing adapters and indexes. The purified PCR products were then pooled and sequenced on an Illumina NovaSeq 6000 platform (S Prime v1.5, 2 x 250 bp).

At each site, a MT was deployed facing south, with collection bottles containing 500 mL of 96% ethanol to preserve captured insects. After one week, the bulk insect samples were retrieved, and six eDNA substrates were collected: water, sediment, spiderwebs, deadwood, anthill material, and soil. All eDNA samples were taken within 10 m of the MT, except for anthill material, which was collected within a 50 m radius due to sparse occurrence, resulting in anthill samples from nine sites.

Water samples were collected following the Complete eDNA Sampling Kit protocol (Sylphium, SYL009-08-20). A total of 300 mL of water was filtered on site through a 0.8 µm polyethersulfone (PEF) membrane filter from three points spaced 2-3 m apart, followed by addition of the preservation solution from the sampling kit. Field blanks were added by filtering Milli-Q water on site. Sediment (∼5 g) was collected from the same water sampling points and stored in a 50 mL Falcon tube. Spiderwebs from three understory sites per location were swabbed and stored in a 50 mL Falcon tube. Deadwood samples (∼5 g) were taken from three decaying spruce logs per location and stored in 50 mL Falcon tubes. Anthill surface material was collected in 50 mL Falcon tubes. To assess within-site reproducibility, we collected eight soil samples per location, one beneath each of the eight spruce trees nearest to the MT. For each sample, about 15 mL of surface soil was taken from three points by a tree after removing leaf litter, pooled, and stored in a single 50 mL Falcon tube. All samples were immediately placed in thermal boxes with cooling blocks and transported to the laboratory within two days, where they were stored at -20°C until further processing.

### 2.2 DNA extraction and metabarcoding sequencing

DNA extraction from MT bulk insect samples was performed following a previously established protocol designed to lyse the insect surface rather than homogenizing the entire body, preserving the physical specimens and minimizing bias caused by interspecies differences in body size (Iwaszkiewicz-Eggebrecht et al., 2023). DNA from water filters was extracted using the Environmental DNA Isolation Kit (Sylphium, SYL002/20/000), optimized for DNA purification from eDNA Dual Filter Capsules. DNA from the other substrates was extracted with the E.Z.N.A.® Soil DNA Kit (D5625-01), following the manufacturer’s protocol. Extraction blanks were included for the bulk insect, filter isolation, and soil extraction procedures. All purified DNA samples were stored at -20°C.

Metabarcoding sequencing libraries targeting the insect cytochrome c oxidase subunit (COI) gene were prepared using a two-step PCR method, following a previously established protocol (Iwaszkiewicz-Eggebrecht et al., 2023). In the first PCR, BF3 and BR3 primers were used to amplify a 458 bp fragment of the insect COI gene (Elbrecht & Leese, 2017). The second PCR incorporated Illumina sequencing adapters and sample-specific indexes. Phusion Green High-Fidelity DNA Polymerase (2 U/µL, F534L) was used for all PCR reactions. Negative PCR controls using nuclease-free water as the template were included in every PCR plate. Following amplification, the second-step PCR products were purified using the ProNex® Size-Selective Purification System, quantified by Qubit 4 Fluorometer (Invitrogen™ dsDNA Quantification Assay Kit), and pooled equimolarly. Sequencing was performed on two NovaSeq 6000 lanes (S Prime v1.5, 2 x 250 bp).

### 2.3 Bioinformatic analysis

Raw reads were demultiplexed using sample indexes. Bioinformatic analysis followed the Qiime*2* v.2024.2 pipeline (Bokulich et al., 2018) run on Linux using the UPPMAX computing cluster. First, the sequencing reads from all samples (excluding blanks) were imported into a single qza format file using Qiime tools import. Adapter trimming was performed to remove primer sequences with qiime cutadapt trim-paired, followed by sequence merging, quality filtering, and denoising using qiime dada2 denoise-paired. Feature tables and representative sequences were generated using qiime feature-table summarize, incorporating sample metadata. Low-frequency ASVs with less than 10 reads across all samples were discarded as potential artefacts.

Prior to taxonomic classification, ASVs were screened using MMSeqs2 v.15-6f452 taxonomy module against the NCBI NR database (February 2024) with default settings except ‘-s 1 –max-seqs 100 --lca-ranks superkingdom,kingdom,phylum,class --tax-lineage 1’, followed by removal of ASVs assigned to bacteria and archaea (Mirdita et al., 2021). Taxonomic assignment of the remaining ASVs followed the approach developed in the Insect Biome Atlas project (Miraldo et al., 2025), combining SINTAX (Edgar, 2016) and EPA-NG v.0.3.8 (Barbera et al., 2019). Specifically, ASVs were initially annotated using SINTAX (implemented in VSEARCH v.2.29.1) against a custom-cleaned reference COI database from BOLD (https://doi.org/10.17044/scilifelab.20514192.v4). Taxonomic assignments were subsequently refined using phylogenetic methods by EPA-NG. The resulting taxonomy file was filtered to retain only insect sequences, and both the feature table and representative sequences were updated accordingly.

The remaining ASVs were clustered into OTUs as part of the HAPP workflow using Swarm with a clustering distance threshold of d = 15. OTU clustering was performed to facilitate comparisons among highly heterogeneous environmental substrates while reducing inflation of diversity estimates caused by sequencing artefacts, residual errors, and intra-specific sequence variation. Sequencing depth was evaluated using per-sample read distributions and rarefaction curves. Rarefaction curves approached asymptotic richness estimates for the majority of samples at approximately 40,000 reads, indicating that additional sequencing would yield relatively few additional OTUs. Based on these results, samples were rarefied to 40,000 reads to standardize sequencing effort across substrates while retaining most observed diversity. Samples with fewer than 40,000 reads were excluded, resulting in the removal of 167 of the 586 samples. The rarefied OTU table was subsequently used for calculating OTU richness, Shannon diversity, Simpson diversity, and Bray-Curtis dissimilarities.

### 2.4 Statistical analysis

Statistical analyses were conducted in R v4.5.0, utilizing the taxonomy file, feature table, and metadata. The phyloseq v.1.48.0 and vegan v.2.6-6.1 packages were used for analysis (Dixon, 2003; McMurdie & Holmes, 2013), while figures were generated with ggplot2 v.3.5.1 and ggpubr v.0.6.0 (Ginestet, 2011).

To evaluate the efficiency of COI metabarcoding for insect monitoring across the seven substrate types, we first calculated IRR and IOR for each sample, followed by a Kruskal-Wallis test to examine their differences between substrates separately. This method was selected because it is robust to non-normal distributions and unequal variances. Post-hoc pairwise comparisons were performed using Dunn’s test with Benjamini-Hochberg (BH) adjustment for multiple comparisons to identify significant pairwise differences.

Insect diversity across substrates and sampling locations was examined using Shannon and Simpson diversity indices, which were calculated using the matrix of relative OTU abundances. We analyzed each index using a Tweedie generalized additive model (GAM) with a log link function, implemented in the package mgcv (v.1.9-1). The model treated substrate type as a fixed factor (with default treatment contrasts) and sampling location as a random intercept using a random-effect smooth term (bs=“re”). Model fitting used restricted maximum likelihood (REML), with significance assessed via approximate p-values. All analyses retained the original zero values of diversity indices (27.5% of samples) through the ability of the Tweedie distribution to accommodate exact zero values. Additionally, a Kruskal-Wallis test was conducted to assess the differences among substrates, followed by post-hoc pairwise comparisons using Dunn’s test with BH adjustment to identify significant pairwise differences.

Analyses of insect composition (beta diversity) were performed using Bray-Curtis dissimilarity based on OTU relative abundance data, excluding samples with zero insects. To assess the effects of both substrate type and sampling location, and their interaction, a PERMANOVA with 999 permutations based on the dissimilarity matrix was conducted using adonis2 in package vegan. Pairwise PERMANOVA analyses were subsequently performed to explore the differences between pairs of substrates using pairwise.adonis2 in package pairwiseAdonis (v0.4.1). Visualization of differences in insect community composition across substrates was conducted using PCoA based on Bray-Curtis dissimilarity. Additionally, we calculated multivariate dispersion (distance-to-centroid) for each substrate to quantify within-substrate heterogeneity using the betadisper function in package vegan. Differences in dispersion among substrates were assessed using the Kruskal-Wallis test, followed by Dunn’s post-hoc tests with BH correction for multiple comparisons, as dispersion values violated assumptions of homogeneity of variance (Levene’s test, p < 0.05). To assess the robustness of the results to the choice of beta-diversity metric, we additionally performed ordination and PERMANOVA analyses using Aitchison distances derived from centered log-ratio (CLR) transformed OTU richness.

Additionally, insect taxonomic composition between substrates was analyzed incorporating different aspects. First, we analyzed insect order composition using both relative OTU richness and relative read abundance data. PERMANOVA with 999 permutations was conducted to test the effects of substrate, location, and their interaction on community composition, followed by pairwise comparisons with BH between substrates using pairwiseAdonis package. PCoA based on Bray-Curtis dissimilarities was employed to visualize compositional differences between substrates. Within-substrate variation was quantified by calculating the average distance to group centroids in multivariate space. The statistical significance of dispersion differences among substrates was assessed using Kruskal-Wallis tests followed by Dunn’s post-hoc pairwise comparisons. Second, the proportion of known vs. unknown taxa (hidden diversity) was evaluated by calculating the percentage of OTUs assignable at each taxonomic level (family, genus, species). A linear mixed-effects model was applied to test the effect of substrate type and taxonomic level, and their interaction on the proportion of known taxa using package lmerTest (v3.1-3). Taxonomic resolution across substrates was assessed by comparing assignment proportions at different taxonomic levels, with statistical significance determined using Kruskal-Wallis tests followed by Dunn’s post-hoc comparisons (Benjamini-Hochberg corrected p-values). Last, the shared and the unique species between substrates were evaluated in more detail.

## 3 Results

To assess the utility of different sampling substrates for eDNA-based insect monitoring, we collected one MT sample and six different types of eDNA substrate samples from each of the 45 sampling locations during autumn (Figure 1). At each site, we collected and sequenced one sample of each substrate type (if available), with the exception of anthill samples (two at one site) and soil samples (eight per site). In some cases, not all sample types could be detected at a given site, in particular anthill samples, which were only found at nine locations (with two samples collected at location 10C), and spiderweb samples, which were collected at 39 locations (Figure 1). Altogether, this resulted in 586 samples subjected to metabarcoding sequencing, yielding 93,381 amplicon sequence variants (ASVs). After data processing and clustering, these ASVs were grouped into 43,620 OTUs, of which 2,714 ASVs corresponding to 937 OTUs were identified as being of insect origin.

### 3.1 Recovery efficiency differs among substrates

To compare the insect-DNA yield among substrates, we calculated the insect read ratio (IRR), defined as the proportion of insect reads relative to the total reads, and the insect OTU ratio (IOR), defined as the number of detected insect OTUs per million sequencing reads. Samples that did not contain any insect OTUs were excluded. Both IRR and IOR differed significantly among substrates (Kruskal-Wallis, p < 0.01). MT samples presented the highest mean IRR (45.40%), whereas eDNA substrates exhibited much lower values, ranging from 0.02% in anthill samples to 2.88% in water samples (Figure 2a). In contrast, water samples had the highest mean IOR (305 OTUs per million reads), followed by MT samples (119 OTUs per million reads) (Figure 2b). Pairwise comparisons indicated that MT and water samples were similar in IOR, while both differed from the other substrates (Table S2). Notably, no insect OTUs were detected in 64 soil and two spiderweb samples.

**Figure 2.**
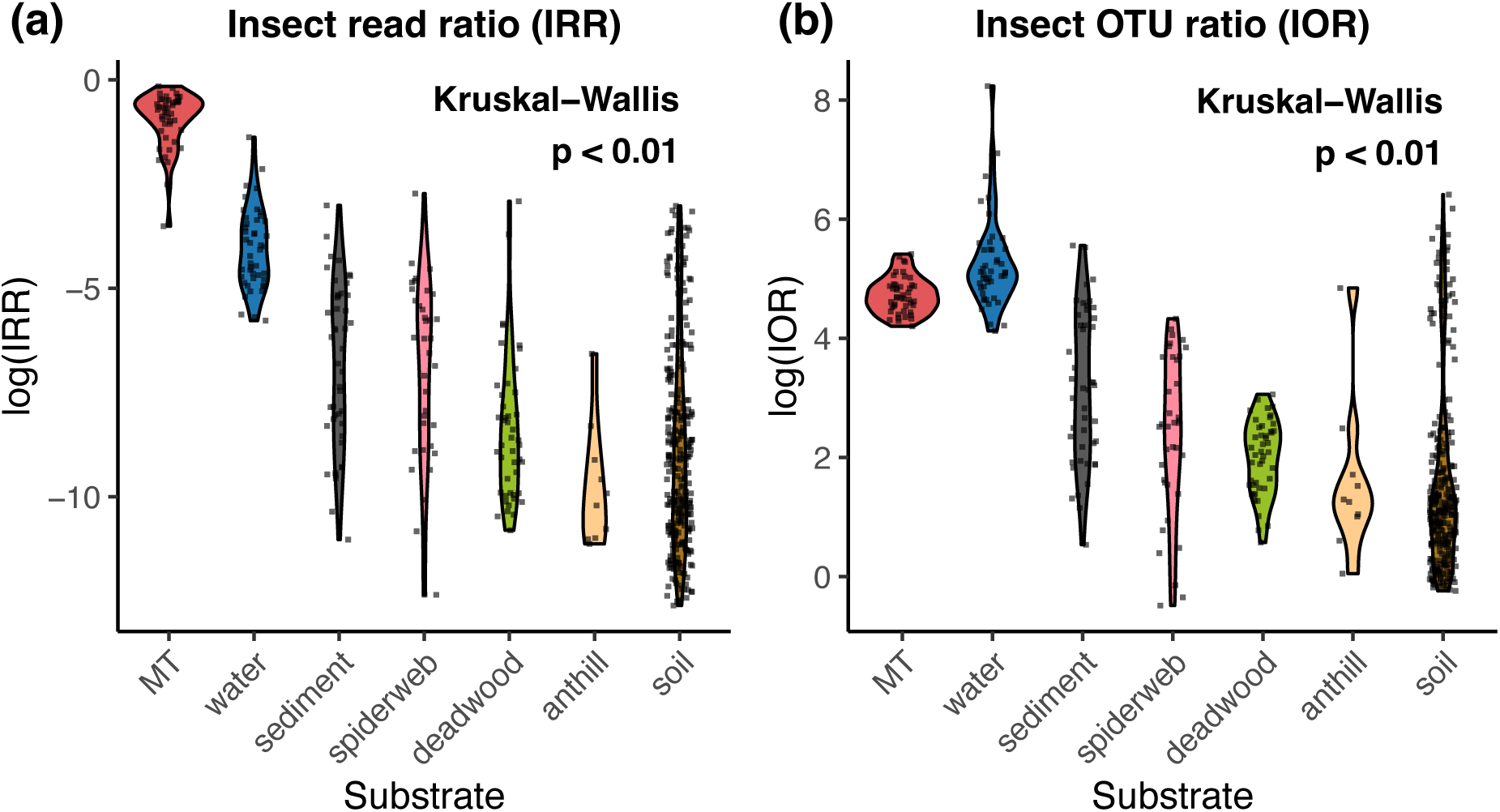
Efficiency of COI metabarcoding for insect DNA detection across seven types of substrates. **(a)** Insect read ratio (IRR), defined as the proportion of insect reads relative to total reads, and **(b)** insect OTU ratio (IOR) defined as the number of insect OTUs per million sequencing reads. Samples with no detected insect OTUs have been removed from this plot.

After quality filtering and rarefaction to 40,000 reads per sample, 419 samples remained, yielding 878 insect OTUs representing 466 species from 16 insect orders (Data S1, Figure S1). The yield of OTUs differed greatly between substrate types, with the mean number of insect OTUs across sites varying between 99 for MTs and 3 for anthills (Figure 3). Spatial patterns were also observed across the sampling sites, where sediment samples show higher insect OTU richness in the northern-most sites (site-clusters 1-6), whereas soil and spiderwebs showed generally higher counts in the southernmost sites (site-clusters 14+15). Overall, OTU richness pooled across all substrates showed a mean number of 165 insect OTUs per site. These pooled OTU richness stayed fairly consistent across sites, with sites in the center of our sampling range (site-clusters 7-11) showing slightly reduced counts, while the counts at the southernmost sites (site-clusters 14+15) are slightly elevated.

**Figure 3.**
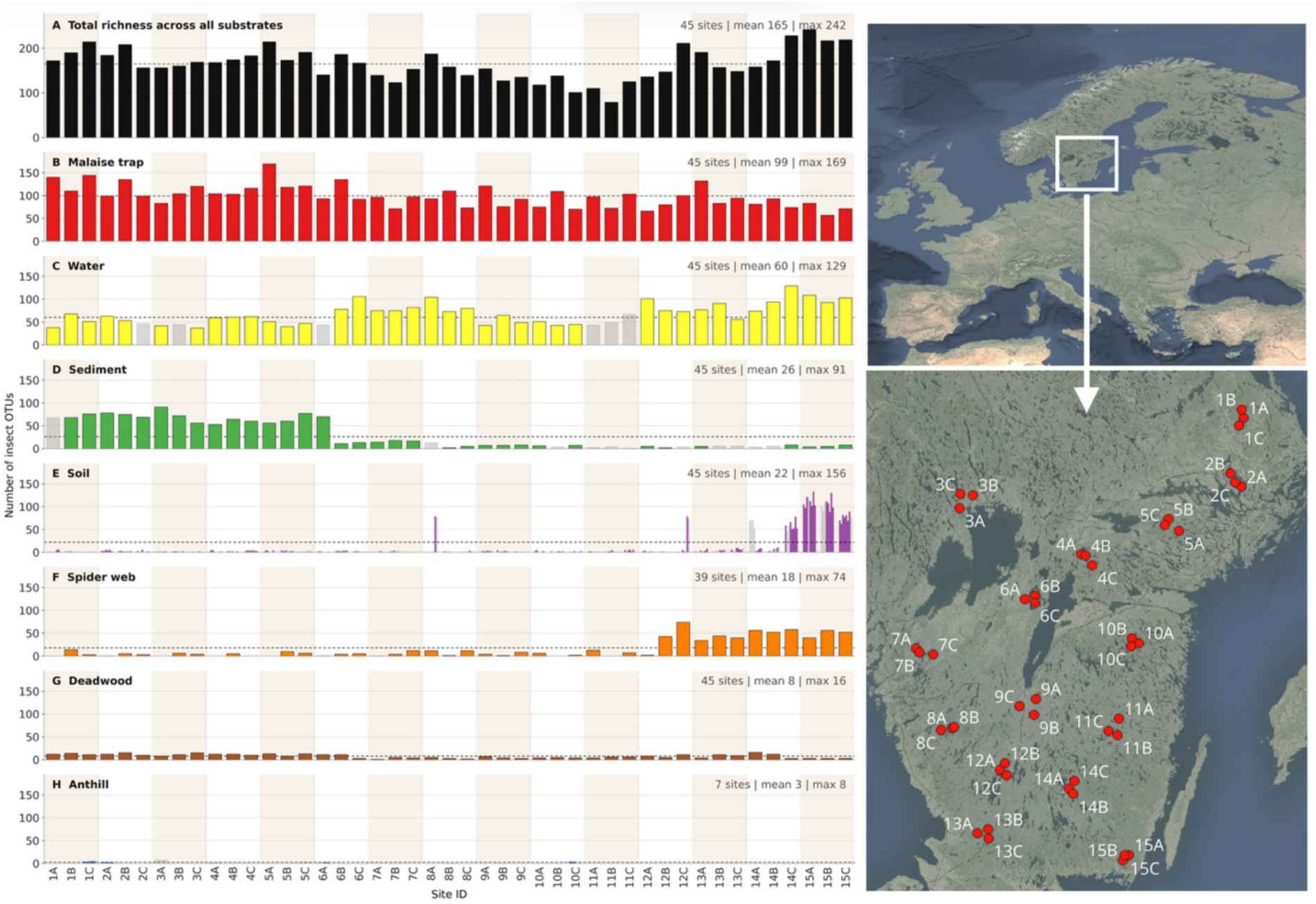
Number of detected insect OTUs per substrate. In case of soil and anthill samples, multiple bars are shown per site, reflecting the multiple subsamples taken per site for these substrates. Samples have been rarefied to 40,000 reads, with samples not clearing that read-threshold shown as grey bars. The height of the grey bars is based on the total number of OTUs detected for these samples using all available reads. The order of site indices on the x-axis approximately follows a north-south gradient (see map panel).

### 3.2 Diversity pattern differs across substrates and locations

To evaluate the effects of substrate type and sampling location on insect diversity, we calculated the OTU richness, Shannon and Simpson diversity indices for each sample (Figure 4a, b and c). A generalized additive model (GAM) revealed highly significant effects of both substrate (F = 35.6, p < 0.01) and location (F = 2.38, p < 0.01) on Shannon diversity, with substrate explaining a substantially greater proportion of the variance. This effect was further supported by a Kruskal-Wallis test (*χ*^2^ = 160.08, p < 0.01), followed by pairwise comparisons of substrate types, showing that over half of the pairs (16 in 21) differed significantly (Table S3). Specifically, water samples exhibited significantly higher Shannon diversity than most other substrates (p < 0.01), except MTs (p > 0.1). Soil samples showed no significant difference from anthill samples (p > 0.1) but were significantly lower than other substrates (p < 0.01, Figure 4b). The Simpson index revealed a broadly similar pattern across substrates and locations, although with generally lower significance levels (Figure 4b, Table S3).

**Figure 4.**
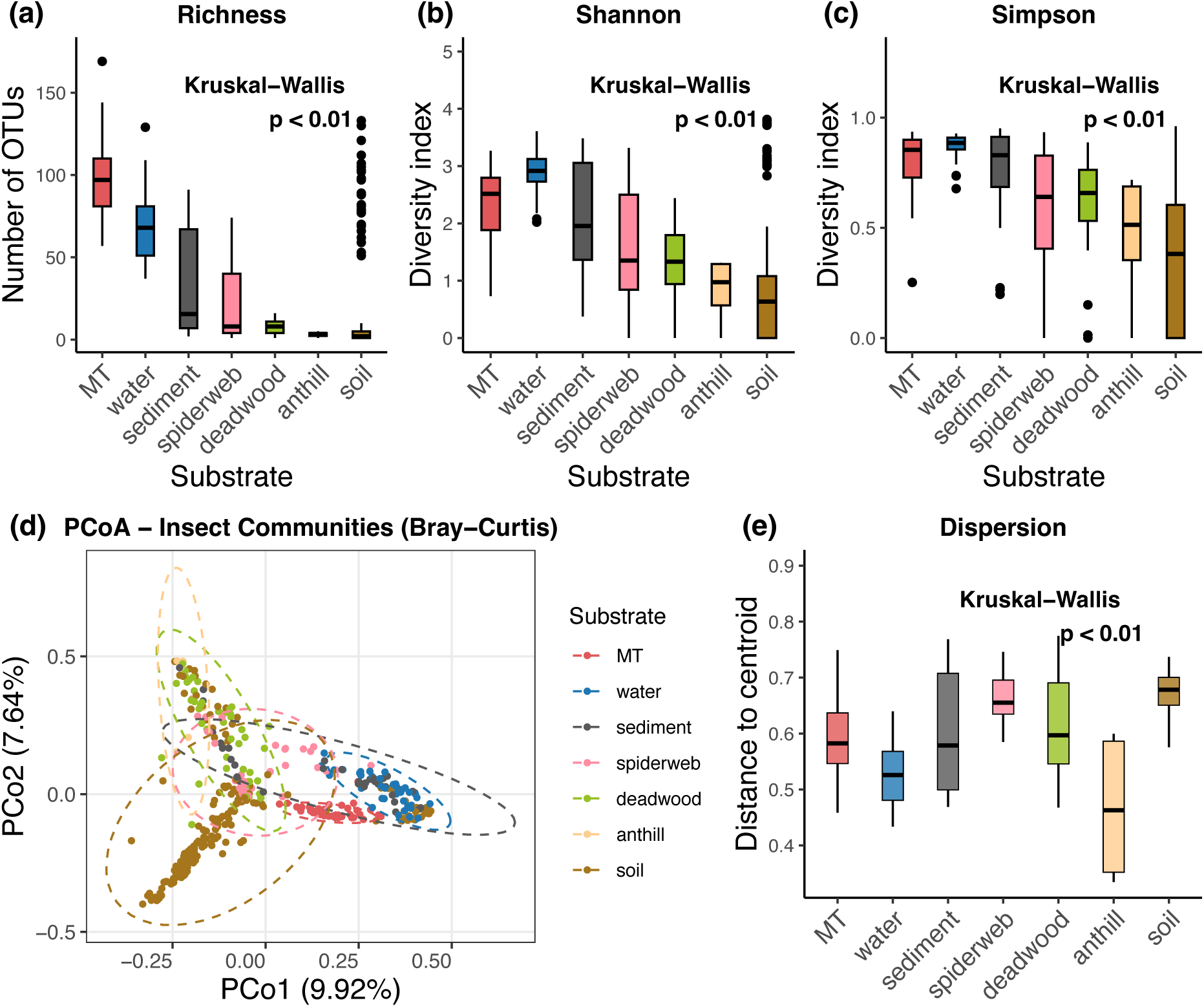
Insect diversity across substrates. Panels **(a), (b)** and **(c)** show Richness, Shannon and Simpson diversity indices calculated from OTU relative abundance data for each substrate. Boxplots show the median (horizontal line), 25th and 75th percentiles. Soil samples showed a multimodal distribution, particularly in regard to OTU richness, with most samples yielding a very low OTU count (< 5) while others showed higher values between 50-150 insect OTUs. **(d)** Principal Coordinates Analysis (PCoA) ordination of Bray-Curtis dissimilarities visualizing beta-diversity patterns among substrates. Points represent individual samples, colored by substrate; ellipses indicate 95% confidence intervals around the centroids of each substrate. **(e)** Dispersion of beta-diversity within substrates, quantified as the distance of samples to their group centroid. Boxplots show the median (horizontal line), 25th and 75th percentiles.

We quantified between-sample differences in insect community composition (beta-diversity) using Bray-Curtis dissimilarities calculated from relative OTU abundances. To test whether compositional dissimilarities were structured by substrate type, sampling location, and their interaction, we used PERMANOVA. Both substrate (R^2^ = 0.12, F = 11.60, p < 0.01) and location (R^2^ = 0.16, F = 2.06, p < 0.01) were significant, and the significant substrate × location interaction (R^2^ = 0.43, F = 1.27, p < 0.01) indicated that differences among substrates were not consistent across locations. Together, these factors explained 70% of the variance in beta diversity. Pairwise PERMANOVA comparisons further confirmed significant differences between all substrate pairs (p < 0.05). Analyses based on Aitchison distances yielded qualitatively similar ordination patterns and identified the same significant effects of substrate type, sampling location, and their interaction (Figure S2).

To visualize these differences, we performed Principal Coordinates Analysis (PCoA) based on the Bray-Curtis dissimilarity matrix (Figure 4d). The ordination plot revealed overall clustering by substrate, with substantial overlap between groups, except soil which shows a more unique community composition. Ellipses indicating 95% of the samples around each substrate centroid were included to illustrate group-level dispersion. Finally, we assessed within-substrate variability by calculating the dispersion of samples around their group centroids (Figure 4e). The degree of dispersion differed significantly among substrates (Kruskal-Wallis test, p < 0.01). MT, sediment, deadwood, and soil samples showed significantly higher dispersion compared to water, spiderweb, and anthill samples (Dunn’s post-hoc tests, p < 0.01, Table S4), highlighting greater community variability within those substrates.

### 3.3 Community composition differs among substrates and locations

Following taxonomic assignment, OTUs were aggregated at the insect order level to characterize taxonomic composition across substrates and locations. We first summarized the total number of OTUs assigned to each insect order within each substrate by pooling all samples (Figure 5a). To account for differences in sample representation among substrates and locations, we additionally calculated the mean relative OTU richness of each insect order across samples within each substrate (Figure 5b) and location (Figure 5c). Diptera was the dominant order across all substrates, accounting for the highest mean relative OTU richness in water samples (77.66%) and the lowest in soil samples (45.34%). Several orders exhibited strong substrate specificity. Odonata was detected only in water samples (0.26% mean relative OTU richness), Raphidioptera only in deadwood samples (0.44%), and Embioptera only in sediment samples (2.97%).

**Figure 5.**
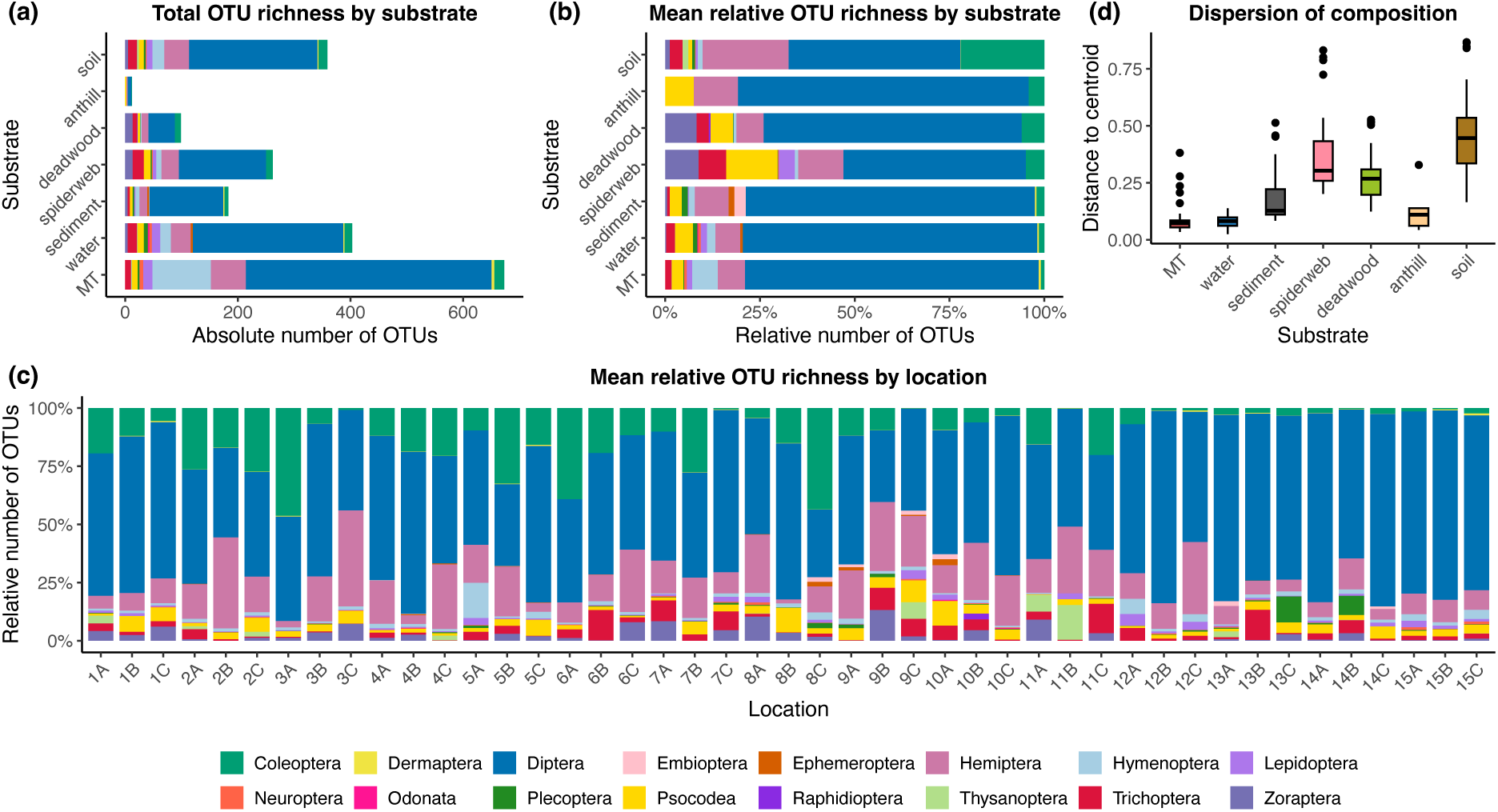
Insect order composition and community dispersion across substrates and sampling locations. Relative OTU richness was calculated for each sample as the proportion of OTUs assigned to each insect order. **(a)** Total OTU richness assigned to each insect order across substrate types, calculated after pooling all samples within each substrate. **(b)** Mean relative OTU richness of insect orders across substrate types. **(c)** Mean relative OTU richness of insect orders across sampling locations. **(d)** Dispersion of insect order composition within each substrate type, measured as the distance of samples to their substrate centroid in multivariate ordination space. Boxplots displayed the median (horizontal line), interquartile range (25th and 75th percentiles), and variation among samples. Colors represent different insect orders in panels **(a)**-**(c)**.

To evaluate differences in insect order composition between substrates and locations, we applied PERMANOVA based on Bray-Curtis dissimilarities calculated from sample-wise relative proportions of OTUs aggregated at the insect order level. Both substrate (R^2^ = 17.49%, F = 13.80, p < 0.01) and location (R^2^ = 18.30%, F = 1.97, p < 0.01) had significant effects on insect community composition, while their interaction was not significant (F < 1). Pairwise PERMANOVA further confirmed significant differences between most substrate pairs (p < 0.05; Table S5), except that anthill samples were only significantly different from water (p < 0.01). PCoA ordination revealed some overlaps between groups (Figure S3). Dispersion to the centroid of each substrate, representing the within substrate variation in order-composition, varied significantly between substrates (Kruskal-Wallis, p < 0.01), with Dunn’s post-hoc tests showing specific pairwise differences (Table S6; Figure 5d).

An analysis based on relative read abundance revealed similar insect order composition pattern to relative OTU abundance (Figure S4; Table S7), both substrate and location showed significant impacts (PERMANOVA, p < 0.01). Diptera contributed higher percentages (51.20-97.26%), compared to 45.34-77.66% based on relative OTU richness (Table S8).

### 3.4 Taxonomic resolution and species recovery across substrates

Among the 878 OTUs after rarefying, 84.74% were classified to family level (109 families), 73.92% to genus (299 genera), and 53.76% to species (466 species). MT samples exhibited the highest taxonomic richness, detecting 86 families and 390 species, followed by water samples with 75 families and 257 species (Table 1). In contrast, soil samples yielded only 215 species despite comprising many more replicates, while anthill samples had the lowest counts overall (8 families and 8 species).

**Table 1.**
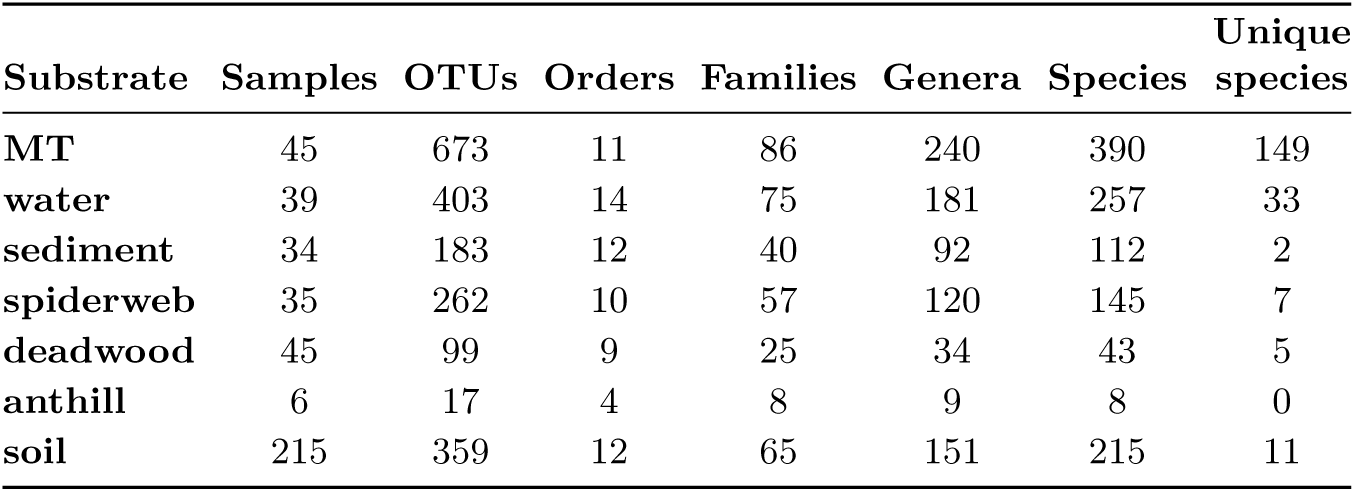
Insect OTU richness and taxonomic summary statistics across substrates. Unique species represents the number of species only found by the substrate.

| Substrate | Samples | OTUs | Orders | Families | Genera | Species | Unique species |
| --- | --- | --- | --- | --- | --- | --- | --- |
| MT | 45 | 673 | 11 | 86 | 240 | 390 | 149 |
| water | 39 | 403 | 14 | 75 | 181 | 257 | 33 |
| sediment | 34 | 183 | 12 | 40 | 92 | 112 | 2 |
| spiderweb | 35 | 262 | 10 | 57 | 120 | 145 | 7 |
| deadwood | 45 | 99 | 9 | 25 | 34 | 43 | 5 |
| anthill | 6 | 17 | 4 | 8 | 9 | 8 | 0 |
| soil | 215 | 359 | 12 | 65 | 151 | 215 | 11 |

The proportion of OTUs successfully assigned to family, genus, and species levels differed only minimally among substrates. The species-level results reveal that a substantial proportion of OTUs for each substrate (44-61%) could not be assigned to a matching species in the reference database, constituting previously unmapped sequence variation (Table 2). Across substrates, over 82% of OTUs were typically assigned to family level except for deadwood (61%). The distribution of assignment proportions for each sample across taxonomic ranks is shown in Figure 6a. A linear mixed-effects model indicated significant main effects of substrate (F = 7.26, p < 0.01) and taxonomic level (F = 93.99, p < 0.01) on the proportion of assigned taxa, as well as a significant interaction (F = 6.22, p < 0.01), suggesting that substrate effects on assignment proportions differed by taxonomic rank. Kruskal–Wallis tests further confirmed significant differences in assignment proportions between substrates at all ranks (p < 0.01), supported by consistent patterns in pairwise comparisons (Table S9).

**Figure 6.**
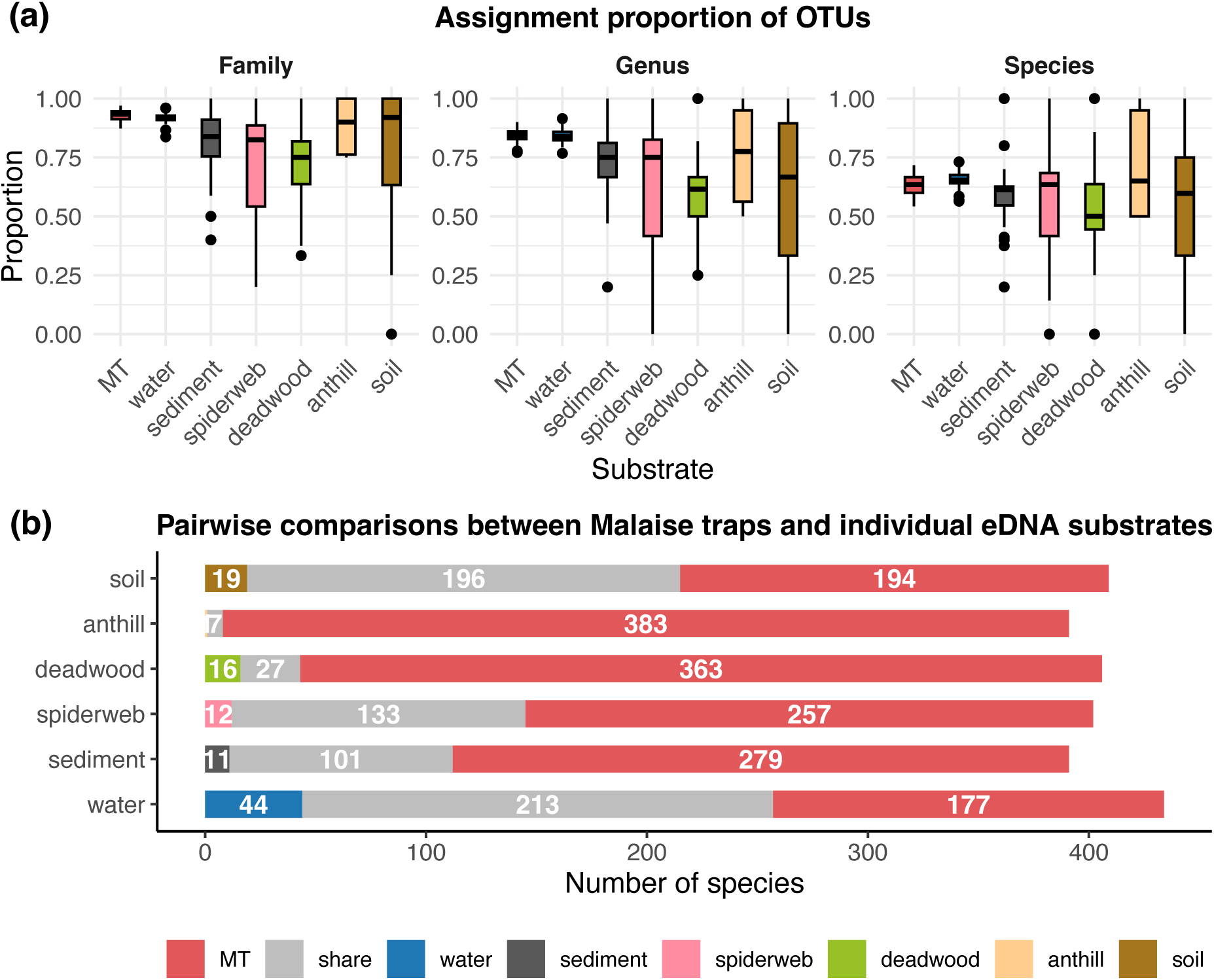
Taxonomic resolution and species recovery across substrates. **(a)** Proportion of insect OTUs successfully assigned to family, genus, and species levels across substrate types. Colors represent different substrates. **(b)** Pairwise species overlap between Malaise traps (MTs) and individual eDNA substrates. Horizontal stacked bars indicate the numbers of species detected exclusively by MTs (MT only), shared between MTs and each eDNA substrate (Shared), or detected exclusively by the corresponding eDNA substrate (eDNA only).

**Table 2.** Number and proportion of OTUs assigned to insect family, genus and species levels across substrates.

| Substrate | family |  | genus |  | species |  |
| --- | --- | --- | --- | --- | --- | --- |
|  | number | proportion | number | proportion | number | proportion |
| MT | 618 | 0.93 | 544 | 0.81 | 391 | 0.58 |
| water | 365 | 0.91 | 336 | 0.83 | 247 | 0.61 |
| sediment | 161 | 0.88 | 143 | 0.78 | 108 | 0.59 |
| spiderweb | 207 | 0.79 | 188 | 0.72 | 142 | 0.54 |
| deadwood | 60 | 0.61 | 49 | 0.50 | 44 | 0.44 |
| anthill | 10 | 0.83 | 8 | 0.67 | 7 | 0.58 |
| soil | 309 | 0.86 | 278 | 0.78 | 207 | 0.58 |

We further examined species overlap and substrate-specific species recovery. Only six species were detected across all substrate types, including five Diptera (*Cheilotrichia cinerascens*, *Cordyla brevicornis*, *Cratyna brevispina*, *Dolichosciara flavipes*, *Scatopsciara atomaria*), and one Psocodea (*Loensia fasciata*), indicating limited species overlap among substrates. Pairwise comparisons between MTs and individual eDNA substrates revealed substantial variation in species overlap (Figure 6b). Water samples exhibited the greatest overlap with MTs and recovered the highest number of species not detected by trapping, whereas sediment, soil, deadwood, and anthill samples contributed comparatively few additional species. Lists of substrate-specific species are provided in Data S2. MTs recovered the highest number of unique species overall (149 species), with Diptera accounting for 94 of these detections (Table S10). Although eDNA substrates generally contributed fewer unique species than MTs, they recovered taxa that were not detected by trapping. Water samples provided the largest contribution of additional species, including 15 unique Diptera species and two Odonata species detected exclusively in this substrate. Spiderweb and deadwood samples primarily contributed unique Coleoptera species, while soil and sediment samples yielded relatively few substrate-specific detections. The number of species shared among substrate pairs is presented in Figure S5.

## 4 Discussion

eDNA metabarcoding has rapidly expanded the range of substrates available for terrestrial biodiversity assessment, yet whether different substrates provide equivalent or complementary representations of ecological communities has remained largely unresolved (Chimeno et al., 2022; Kirse et al., 2021; Valentin et al., 2020). By comparing MT and six eDNA substrates across 45 locations using a standardized sampling and analytical framework, our study demonstrates that substrates capture complementary rather than equivalent representations of insect biodiversity. This conclusion is consistently supported by differences in insect detection efficiency, diversity, community composition, taxonomic representation, and species recovery across substrates. Although MTs recovered the greatest overall richness, no single substrate represented the full spectrum of insect biodiversity, and each substrate contributed unique biodiversity information that was absent from others. Together, these findings indicate that substrate choice influences not only how many taxa are detected but also which components of insect communities become represented in metabarcoding datasets. Instead of serving as interchangeable sampling media, environmental substrates provide complementary representations of terrestrial insect biodiversity, with important consequences for both ecological interpretation and biodiversity monitoring.

The complementary biodiversity patterns observed among substrates are consistent with the distinct ecological processes that govern the production, transport, accumulation, and persistence of eDNA (Newton et al., 2025; Zhao & Andermann, 2026). Rather than representing a homogeneous pool of biological material, terrestrial eDNA is continuously redistributed through multiple pathways before being retained within different environmental substrates (Bell et al., 2024). Consequently, each substrate samples only a subset of the DNA circulating through an ecosystem, and the recovered community reflects both the underlying insect assemblage and the processes controlling DNA movement and preservation. Differences among substrates therefore arise not simply from variation in detection efficiency, but from differences in the ecological information that each substrate is able to retain. Viewed from this perspective, environmental substrates represent complementary observational windows into terrestrial insect communities rather than alternative measurements of the same biodiversity signal.

The contrasting biodiversity patterns recovered from different substrates are consistent with this interpretation. Flowing water integrates DNA originating from both aquatic and surrounding terrestrial habitats through hydrological connectivity, producing spatially integrated biodiversity signals that explain its consistently high OTU recovery and substantial contribution of additional taxa (Stothut et al., 2024; Takenaka et al., 2024). In contrast, soils predominantly retain DNA close to its source, where fine-scale variation in microhabitat conditions, microbial activity, and substrate properties likely contributes to the pronounced spatial heterogeneity observed among samples (Ariza et al., 2023; Hermans et al., 2022; Kong et al., 2026). Spiderwebs intercept airborne organisms, body fragments, and environmental particles moving through the local environment, recovering insect assemblages that differed markedly from those detected in other substrates (Gregorič et al., 2022; Newton et al., 2024). Similar ecological processes are likely to influence sediment (Sakata et al., 2020), deadwood (Purahong et al., 2019), and other environmental substrates, each preserving different subsets of biodiversity information. Together, these observations suggest that complementary biodiversity patterns emerge because different substrates archive eDNA through different ecological pathways, rather than because one substrate universally outperforms another.

Accurate ecological inference depends on understanding what eDNA observations actually represent (Çevik & Çevik, 2025; Ruppert et al., 2019). Our results indicate that terrestrial eDNA datasets derived from different substrates should not be interpreted as equivalent representations of local insect communities, because the recovered assemblages reflect both underlying community composition and the ecological processes governing DNA deposition, transport, accumulation, and persistence within each substrate (Jo et al., 2025; Valentin et al., 2021). Consequently, comparisons among terrestrial eDNA datasets should account for both underlying ecological variation and the substrate-specific representation of biodiversity inherent to different environmental substrates. Recognizing this distinction shifts substrate identity from a methodological variable to an ecological attribute that shapes the biological information available for downstream analyses. Consequently, substrate identity should be explicitly incorporated into the interpretation of terrestrial eDNA datasets, allowing biologically meaningful differences among substrates to inform, rather than confound, ecological inference.

Although substrate complementarity was consistently observed across our study sites, its magnitude is likely to vary among ecosystems, seasons, and taxonomic groups. Our comparison was conducted in spruce-dominated forests during a single autumn sampling period using a standardized COI metabarcoding workflow. Seasonal turnover in insect communities and eDNA dynamics has been reported in previous studies (Reinholdt Jensen et al., 2021), suggesting that substrate-specific biodiversity patterns may also vary through time. Likewise, substrate performance is expected to depend on habitat context and target taxa, and our limited number of anthill samples restricts inference about their relative contribution. These considerations define the conditions under which our conclusions should be interpreted and extended to other systems. Nevertheless, because all substrates were collected and analyzed within a common experimental framework, the observed differences provide robust evidence that substrate identity is an important determinant of biodiversity patterns recovered from terrestrial eDNA.

Our study demonstrates that eDNA substrates recover complementary rather than equivalent dimensions of insect biodiversity. This complementarity arises because different substrates retain distinct ecological information through the processes governing eDNA deposition, transport, accumulation, and persistence, resulting in different representations of the same biological communities. Consequently, substrate choice should be regarded not simply as a methodological decision but as an ecological consideration that directly influences biodiversity inference. As terrestrial eDNA monitoring continues to expand across ecosystems and taxonomic groups, integrating complementary environmental substrates offers a more complete and biologically informative representation of biodiversity than reliance on any single substrate alone. Recognizing that environmental substrates capture complementary dimensions of biodiversity provides a conceptual basis for designing future terrestrial eDNA surveys and for interpreting eDNA observations across ecological studies.

## Acknowledgments

All computations were carried out on the Uppmax computing cluster, which is part of the National Academic Infrastructure for Super-computing in Sweden (NAISS), financially supported by the Swedish Research Council. Sequencing was performed by the SNP&SEQ Technology Platform in Uppsala. The facility is part of the National Genomics Infrastructure (NGI) Sweden and Science for Life Laboratory (SciLifeLab). The SNP&SEQ Platform is also supported by the Swedish Research Council and the Knut and Alice Wallenberg Foundation. Further, we thank the insect research station Linné on Öland, Sweden, for providing MTs that were needed to carry out the fieldwork for this study.

## Data Accessibility and Benefit-Sharing Section

The insect bulk samples collected from MTs are deposited in the wet entomology collections of the Museum of Evolution, Uppsala University (accession number 2025:024). Metabarcoding sequencing data have been deposited in the European Nucleotide Archive (ENA) under accession number ERP166070: https://www.ebi.ac.uk/ena/browser/view/ERP166070. All data needed to evaluate the conclusions in the paper are present in the paper and/or the Supplementary Materials.

## Author Contributions

B.Z. and T.A. conceived and designed the study. B.Z. conducted field sampling, laboratory work, data analyses, and drafted the manuscript. J.S. contributed to bioinformatic and statistical analyses and reviewed the manuscript. T.A. contributed to data analyses, interpretation of results, and manuscript writing. All authors discussed the results, revised the manuscript, and approved the final version.

## References

Ariza, M., Fouks, B., Mauvisseau, Q., Halvorsen, R., Alsos, I. G., & de Boer, H. J. (2023). Plant biodiversity assessment through soil eDNA reflects temporal and local diversity. Methods in Ecology and Evolution, 14 (2), 415–430. 10.1111/2041-210X.13865

Banerjee, P., Al-Bayer, S., Calaor, J., Weber, S., Graham, N. R., Andersen, J. C., Economo, E. P., Kennedy, S., Krehenwinkel, H., Gillespie, R. G., Roderick, G. K., Rogers, H. S., & Puliafico, K. P. (2026). Comparison of Environmental DNA and Bulk DNA Metabarcoding for Assessing Terrestrial Arthropod Diversity Across Three Habitat Types on Guam. Molecular Ecology Resources, 26 (5), e70172. 10.1111/1755-0998.70172

Barbera, P., Kozlov, A. M., Czech, L., Morel, B., Darriba, D., Flouri, T., & Stamatakis, A. (2019). EPAng: Massively Parallel Evolutionary Placement of Genetic Sequences. Systematic Biology, 68 (2), 365–369. 10.1093/SYSBIO/SYY054

Bell, K. L., Campos, M., Hoffmann, B. D., Encinas-Viso, F., Hunter, G. C., & Webber, B. L. (2024). Environmental DNA methods for biosecurity and invasion biology in terrestrial ecosystems: Progress, pitfalls, and prospects. Science of The Total Environment, 926, 171810. 10.1016/J.SCITOTENV.2024.171810

Bohmann, K., & Lynggaard, C. (2023). Transforming terrestrial biodiversity surveys using airborne eDNA. Trends in Ecology and Evolution, 38 (2), 119–121. 10.1016/j.tree.2022.11.006

Bokulich, N. A., Kaehler, B. D., Rideout, J. R., Dillon, M., Bolyen, E., Knight, R., Huttley, G. A., & Gregory Caporaso, J. (2018). Optimizing taxonomic classification of marker-gene amplicon sequences with QIIME 2’s q2-feature-classifier plugin. Microbiome, 6 (1), 1–17. 10.1186/S40168-018-0470-Z

Çevik, T., & Çevik, N. (2025). Environmental DNA (eDNA): A review of ecosystem biodiversity detection and applications. Biodiversity and Conservation 2025 34:9, 34 (9), 2999–3035. 10.1007/S10531-025-03112-Y

Chimeno, C., Hausmann, A., Schmidt, S., Raupach, M. J., Doczkal, D., Baranov, V., Hübner, J., Höcherl, A., Albrecht, R., Jaschhof, M., Haszprunar, G., & Hebert, P. D. N. (2022). Peering into the Darkness: DNA Barcoding Reveals Surprisingly High Diversity of Unknown Species of Diptera (Insecta) in Germany. Insects 2022, Vol. 13, Page 82, 13 (1), 82. 10.3390/INSECTS13010082

Dixon, P. (2003). VEGAN, a package of R functions for community ecology. Journal of Vegetation Science, 14 (6), 927–930. 10.1111/J.1654-1103.2003.TB02228.X

Edgar, R. C. (2016). SINTAX: a simple non-Bayesian taxonomy classifier for 16S and ITS sequences. BioRxiv, 074161. 10.1101/074161

Egeter, B., Peixoto, S., Brito, J. C., Jarman, S., Puppo, P., & Velo-Antón, G. (2018). Challenges for assessing vertebrate diversity in turbid Saharan water-bodies using environmental DNA. Genome, 61 (11), 807–814. 10.1139/gen-2018-0071

Elbrecht, V., & Leese, F. (2017). Validation and development of COI metabarcoding primers for freshwater macroinvertebrate bioassessment. Frontiers in Environmental Science, 5 (APR), 237020. 10.3389/FENVS.2017.00011

Foucher, A., Evrard, O., Ficetola, G. F., Gielly, L., Poulain, J., Giguet-Covex, C., Laceby, J. P., Salvador-Blanes, S., Cerdan, O., & Poulenard, J. (2020). Persistence of environmental DNA in cultivated soils: implication of this memory effect for reconstructing the dynamics of land use and cover changes. Scientific Reports, 10 (1), 10502. 10.1038/s41598-020-67452-1

Ginestet, C. (2011). ggplot2: Elegant Graphics for Data Analysis. Journal of the Royal Statistical Society: Series A (Statistics in Society), 174 (1), 245–246. 10.1111/j.1467-985X.2010.00676_9.x

Gregorič, M., Kutnjak, D., Bačnik, K., Gostinčar, C., Pecman, A., Ravnikar, M., & Kuntner, M. (2022). Spider webs as eDNA samplers: Biodiversity assessment across the tree of life. Molecular Ecology Resources, 22 (7), 2534–2545. 10.1111/1755-0998.13629

Guthrie, A. M., Nevill, P., Cooper, C. E., Bateman, P. W., & van der Heyde, M. (2023). On a roll: a direct comparison of extraction methods for the recovery of eDNA from roller swabbing of surfaces. BMC Research Notes, 16 (1), 1–6. 10.1186/S13104-023-06669-5/FIGURES/3

Hermans, S. M., Lear, G., Buckley, T. R., & Buckley, H. L. (2022). Environmental DNA sampling detects between-habitat variation in soil arthropod communities, but is a poor indicator of fine-scale spatial and seasonal variation. Ecological Indicators, 140. 10.1016/j.ecolind.2022.109040

Iwaszkiewicz-Eggebrecht, E., Łukasik, P., Buczek, M., Deng, J., Hartop, E. A., Havnås, H., Prus-Frankowska, M., Ugarph, C. R., Viteri, P., Andersson, A. F., Roslin, T., Tack, A. J. M., Ronquist, F., & Miraldo, A. (2023). FAVIS: Fast and versatile protocol for non-destructive metabarcoding of bulk insect samples. PLOS ONE, 18 (7), e0286272. 10.1371/JOURNAL.PONE.0286272

Jo, T. S., Murakami, H., & Nakadai, R. (2025). Spatial dispersal of environmental DNA particles in lentic and marine ecosystems: An overview and synthesis. Ecological Indicators, 174, 113469. 10.1016/J.ECOLIND.2025.113469

Johnson, D. M., & Haynes, K. J. (2023). Spatiotemporal dynamics of forest insect populations under climate change. Current Opinion in Insect Science, 56, 101020. 10.1016/J.COIS.2023.101020

Karlsson, D., Hartop, E., Forshage, M., Jaschhof, M., & Ronquist, F. (2020). The Swedish Malaise Trap Project: A 15 Year Retrospective on a Countrywide Insect Inventory. Biodiversity Data Journal 8: E47255, 8, e47255-. 10.3897/BDJ.8.E47255

Kirse, A., Bourlat, S. J., Langen, K., & Fonseca, V. G. (2021). Metabarcoding Malaise traps and soil eDNA reveals seasonal and local arthropod diversity shifts. Scientific Reports 2021 11:1, 11 (1), 1–12. 10.1038/s41598-021-89950-6

Kong, Y., Tian, J., Chai, Z., Li, S., & Yao, M. (2026). Soil eDNA Biomonitoring: Assessing Efficacy for Detecting Terrestrial Vertebrate and Plant Biodiversity. Environmental Science & Technology, 60 (26), 18707–18722. 10.1021/ACS.EST.6C04825

Lunghi, E., Valle, B., Guerrieri, A., Bonin, A., Cianferoni, F., Manenti, R., & Ficetola, G. F. (2022). Environmental DNA of insects and springtails from caves reveals complex processes of eDNA transfer in soils. Science of the Total Environment, 826. 10.1016/j.scitotenv.2022.154022

Marquina, D., Esparza-Salas, R., Roslin, T., & Ronquist, F. (2019). Establishing arthropod community composition using metabarcoding: Surprising inconsistencies between soil samples and preservative ethanol and homogenate from Malaise trap catches. Molecular Ecology Resources, 19 (6), 1516–1530. 10.1111/1755-0998.13071

May, R. M. (1986). Biological diversity: How many species are there? Nature 1986 324:6097, 324 (6097), 514–515. 10.1038/324514a0

McMurdie, P. J., & Holmes, S. (2013). phyloseq: An R Package for Reproducible Interactive Analysis and Graphics of Microbiome Census Data. PLOS ONE, 8 (4), e61217. 10.1371/JOURNAL.PONE.0061217

Miraldo, A., Sundh, J., Iwaszkiewicz-Eggebrecht, E., Buczek, M., Goodsell, R., Johansson, H., Fisher, B. L., Rahar-injanahary, D., Rajoelison, E. T., Ranaivo, C., Randrianandrasana, C., Rafanomezantsoa, J. J., Manoharan, L., Granqvist, E., van Dijk, L. J. A., Alberg, L., Åhlén, D., Aspebo, M., Åström, S., . . . Ronquist, F. (2025). Data of the Insect Biome Atlas: a metabarcoding survey of the terrestrial arthropods of Sweden and Madagascar. Scientific Data 2025 12:1, 12 (1), 835-. 10.1038/s41597-025-05151-0

Mirdita, M., Steinegger, M., Breitwieser, F., Söding, J., & Levy Karin, E. (2021). Fast and sensitive taxonomic assignment to metagenomic contigs. Bioinformatics, 37 (18), 3029–3031. 10.1093/BIOINFORMATICS/BTAB184

Nagler, M., Podmirseg, S. M., Ascher-Jenull, J., Sint, D., & Traugott, M. (2022). Why eDNA fractions need consideration in biomonitoring. Molecular Ecology Resources. 10.1111/1755-0998.13658

Newton, J. P., Allentoft, M. E., Bateman, P. W., van der Heyde, M., & Nevill, P. (2025). Targeting Terrestrial Vertebrates With eDNA: Trends, Perspectives, and Considerations for Sampling. Environmental DNA, 7 (1), e70056. 10.1002/EDN3.70056

Newton, J. P., Nevill, P., Bateman, P. W., Campbell, M. A., & Allentoft, M. E. (2024). Spider webs capture environmental DNA from terrestrial vertebrates. IScience, 27 (2). 10.1016/j.isci.2024.108904

Outhwaite, C. L., McCann, P., & Newbold, T. (2022). Agriculture and climate change are reshaping insect biodiversity worldwide. Nature 2022 605:7908, 605 (7908), 97–102. 10.1038/s41586-022-04644-x

Purahong, W., Mapook, A., Wu, Y. T., & Chen, C. T. (2019). Characterization of the castanopsis carlesii deadwood mycobiome by pacbio sequencing of the full-length fungal nuclear ribosomal internal transcribed spacer (ITS). Frontiers in Microbiology, 10 (MAY), 426646. 10.3389/FMICB.2019.00983/TEXT

Reinholdt Jensen, M., Egelyng Sigsgaard, E., Agersnap, S., Jessen Rasmussen, J., Baattrup-Pedersen, A., Wiberg-Larsen, P., & Francis Thomsen, P. (2021). Seasonal turnover in community composition of stream-associated macroinvertebrates inferred from freshwater environmental DNA metabarcoding. Environmental DNA, 3 (4), 861–876. 10.1002/EDN3.193

Ruppert, K. M., Kline, R. J., & Rahman, M. S. (2019). Past, present, and future perspectives of environmental DNA (eDNA) metabarcoding: A systematic review in methods, monitoring, and applications of global eDNA. In Global Ecology and Conservation (Vol. 17, p. e00547). Elsevier B.V. 10.1016/j.gecco.2019.e00547

Ryan, E., Bateman, P., Fernandes, K., van der Heyde, M., & Nevill, P. (2022). eDNA metabarcoding of log hollow sediments and soils highlights the importance of substrate type, frequency of sampling and animal size, for vertebrate species detection. Environmental DNA, 4 (4), 940–953. 10.1002/edn3.306

Sakata, M. K., Yamamoto, S., Gotoh, R. O., Miya, M., Yamanaka, H., & Minamoto, T. (2020). Sedimentary eDNA provides different information on timescale and fish species composition compared with aqueous eDNA. Environmental DNA, 2 (4), 505–518. 10.1002/edn3.75

Scudder, G. G. E. (2017). The Importance of Insects. Insect Biodiversity, 9–43. 10.1002/9781118945568.CH2

Shogren, A. J., Tank, J. L., Egan, S. P., August, O., Rosi, E. J., Hanrahan, B. R., Renshaw, M. A., Gantz, C. A., & Bolster, D. (2018). Water Flow and Biofilm Cover Influence Environmental DNA Detection in Recirculating Streams. Environmental Science and Technology, 52 (15), 8530–8537. 10.1021/acs.est.8b01822

Stothut, M., Kühne, D., Ströbele, V., Mahla, L., Künzel, S., & Krehenwinkel, H. (2024). Environmental DNA metabarcoding reliably recovers arthropod interactions which are frequently observed by video recordings of flowers. Environmental DNA, 6 (3), e550. 10.1002/EDN3.550

Takenaka, M., Hasebe, Y., Yano, K., Okamoto, S., Tojo, K., Seki, M., Sekiguchi, S., Jitsumasa, T., Morohashi, N., Handa, Y., & Sakaba, T. (2024). Environmental DNA metabarcoding on aquatic insects: Comparing the primer sets of MtInsects-16S based on the mtDNA 16S and general marker based on the mtDNA COI region. Environmental DNA, 6 (4), e588. 10.1002/EDN3.588

Thomsen, P. F., & Willerslev, E. (2015). Environmental DNA – An emerging tool in conservation for monitoring past and present biodiversity. Biological Conservation, 183, 4–18. 10.1016/J.BIOCON.2014.11.019

Tihelka, E., Cai, C., Giacomelli, M., Lozano-Fernandez, J., Rota-Stabelli, O., Huang, D., Engel, M. S., Donoghue, P. C. J., & Pisani, D. (2021). The evolution of insect biodiversity. Current Biology, 31 (19), R1299–R1311. 10.1016/J.CUB.2021.08.057

Valentin, R. E., Fonseca, D. M., Gable, S., Kyle, K. E., Hamilton, G. C., Nielsen, A. L., & Lockwood, J. L. (2020). Moving eDNA surveys onto land: Strategies for active eDNA aggregation to detect invasive forest insects. Molecular Ecology Resources, 20 (3), 746–755. 10.1111/1755-0998.13151

Valentin, R. E., Kyle, K. E., Allen, M. C., Welbourne, D. J., & Lockwood, J. L. (2021). The state, transport, and fate of aboveground terrestrial arthropod eDNA. Environmental DNA, 3 (6), 1081–1092. 10.1002/edn3.229

van der Heyde, M., Bunce, M., & Nevill, P. (2022). Key factors to consider in the use of environmental DNA metabarcoding to monitor terrestrial ecological restoration. Science of The Total Environment, 848, 157617. 10.1016/J.SCITOTENV.2022.157617

van der Heyde, M., Bunce, M., Wardell-Johnson, G., Fernandes, K., White, N. E., & Nevill, P. (2020). Testing multiple substrates for terrestrial biodiversity monitoring using environmental DNA metabarcoding. Molecular Ecology Resources, 20 (3), 732–745. 10.1111/1755-0998.13148

Wu, S., Wang, Y., Qin, H., Zhang, Z., Liu, S., Ruan, Y., Chen, G., Yuan, X., & Zhang, H. (2026). Environmental DNA (eDNA) Technology in Biodiversity and Ecosystem Health Research: Advances and Prospects. Ecology and Evolution, 16 (1), e72891. 10.1002/ECE3.72891

Yao, M., Zhang, S., Lu, Q., Chen, X., Zhang, S. Y., Kong, Y., & Zhao, J. (2022). Fishing for fish environmental DNA: Ecological applications, methodological considerations, surveying designs, and ways forward. Molecular Ecology, 31 (20), 5132–5164. 10.1111/MEC.16659

Zhao, B., & Andermann, T. (2026). Properties and Limitations of eDNA Substrates for Terrestrial Animal Monitoring. Molecular Ecology Resources, 26 (2), e70096. 10.1111/1755-0998.70096

