## Supplemental Figures S1 to S7 and Tables S1 to S10 for "Environmental DNA substrates capture complementary dimensions of insect biodiversity"

**Contents:**

Figures S1 to S7

Tables S1 to S10

Caption of Data S1 and S2

Figures

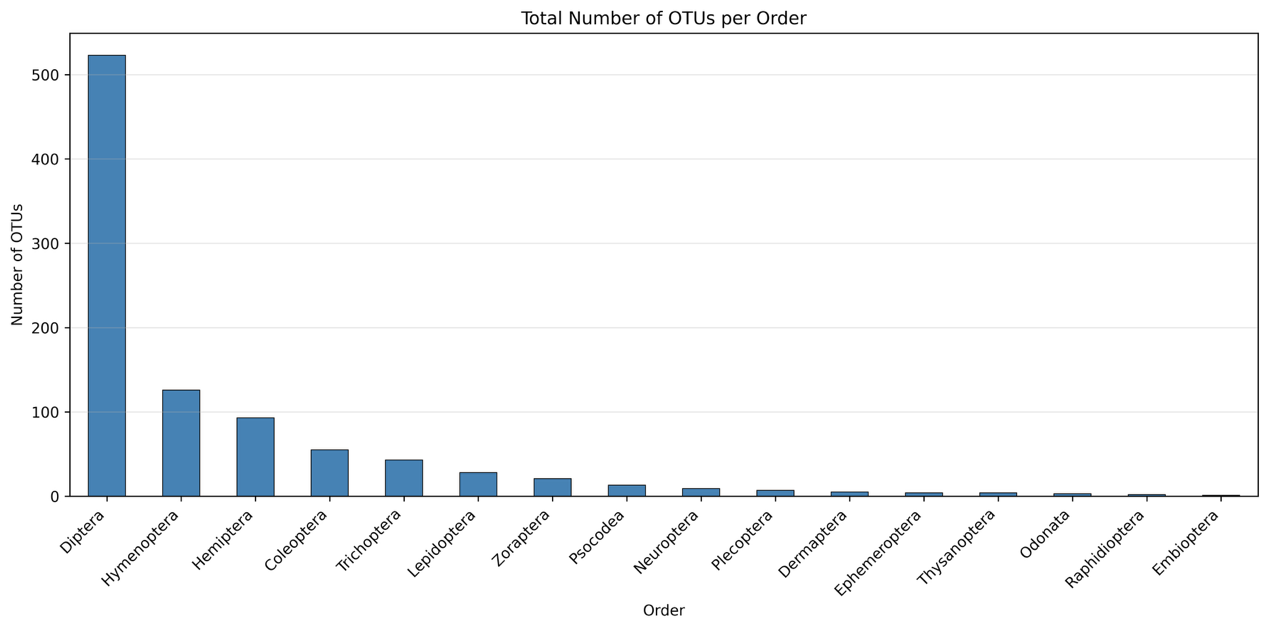

Figure S1. Total number of detected OTUs assigned to each insect order across all samples included in the study. Orders are arranged according to the number of recovered OTUs.

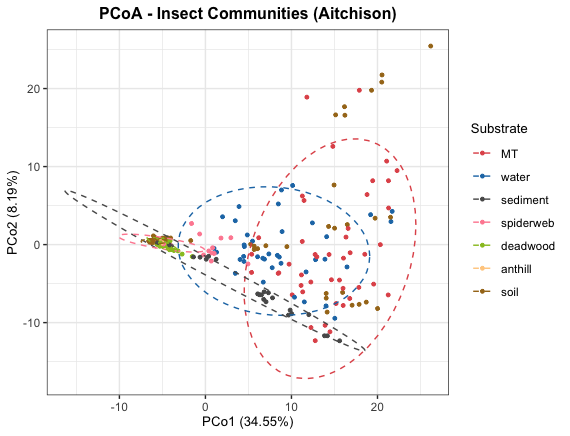

Figure S2. Principal coordinate analysis (PCoA) of insect OTU composition based on Aitchison distances. It shows the distribution patterns of samples across substrates. Each point represents a sample, colored by substrate, and ellipses represent the 95% confidence interval around group centroids.

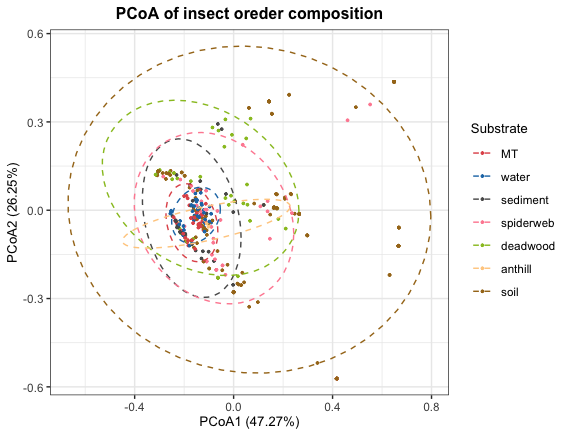

Figure S3. Principal coordinates analysis (PCoA) ordination of insect order composition (using relative OTU abundance data) based on Bray-Curtis dissimilarities. Points represent individual sample, colored by substrate; ellipses indicate 95% confidence intervals around the centroids of each substrate.

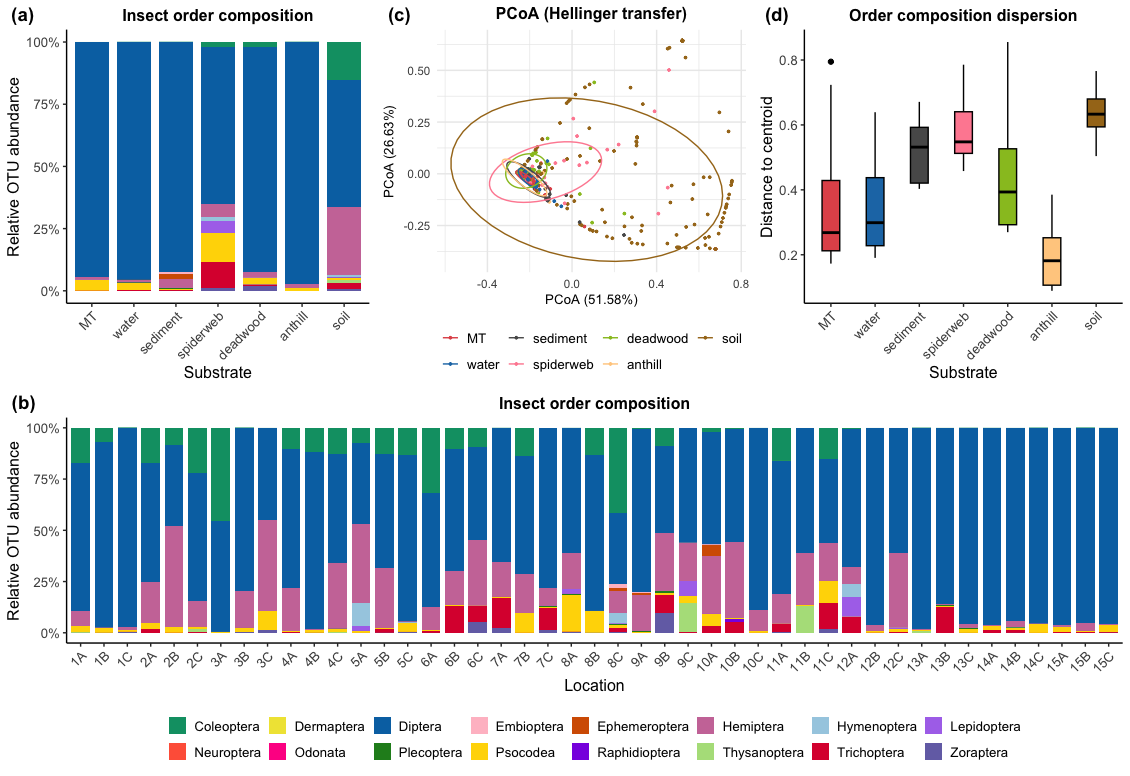

Figure S4. Variation in insect order composition across substrates and locations based on the relative OTU abundance. a Mean relative abundance of insect orders across different substrates. Colors represent different insect orders. b Mean relative abundance of insect orders across different sampling locations. Colors represent different insect orders. c Principal coordinates analysis (PCoA) ordination of insect order composition based on Bray-Curtis dissimilarities, showing clustering of samples by substrates. Each point represents a sample, colored by substrate, and ellipses represent the 80% confidence interval around group centroids. d Dispersion of insect order composition within each substrate group, measured as the distance of samples from their group centroid. Boxplots displayed the median (horizontal line), interquartile range (25th and 75th percentiles), and variation among samples.

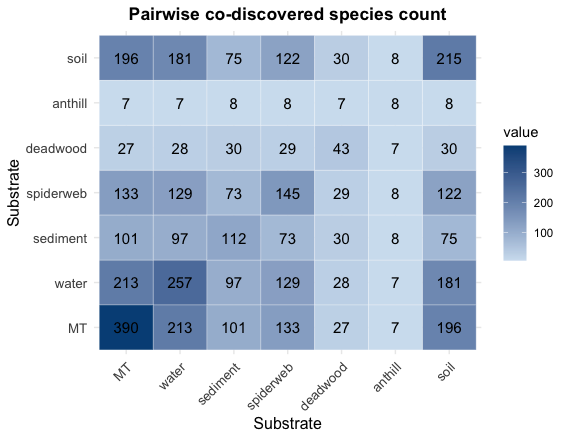

Figure S5. Species counts that were co-discovered between pairs of substrates.

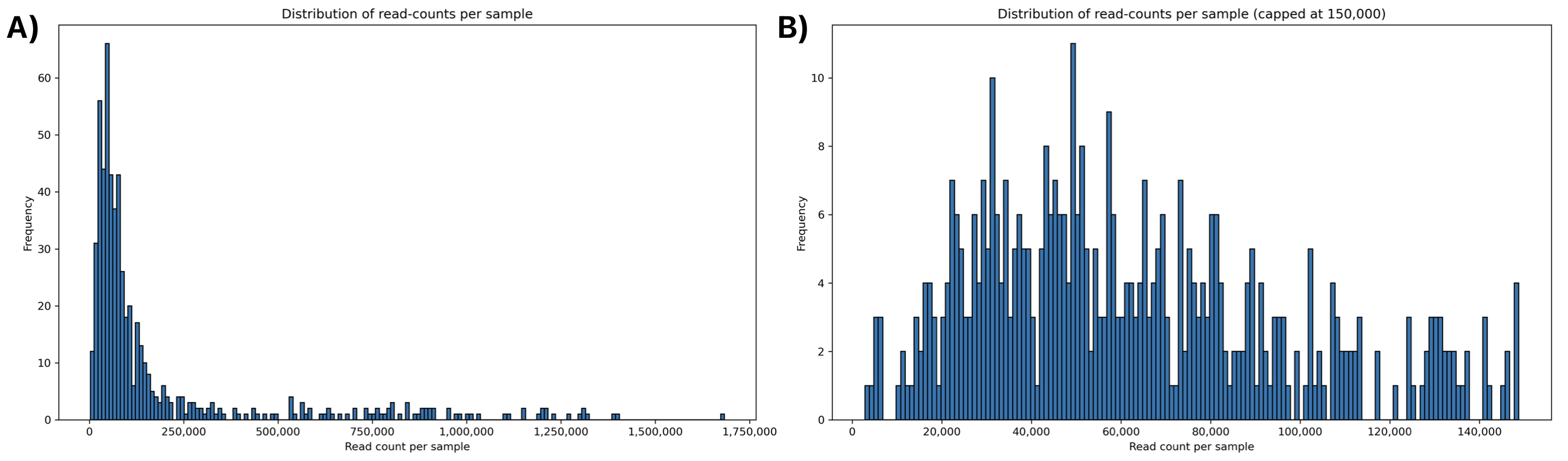

Figure S6. Distribution of sequencing read counts across samples. Panel A displays the full range of read counts across all samples (up to approximately 1,700,000 reads). Panel B provides a zoomed-in view of samples with lower sequencing depth, showing read counts up to 150,000 reads.

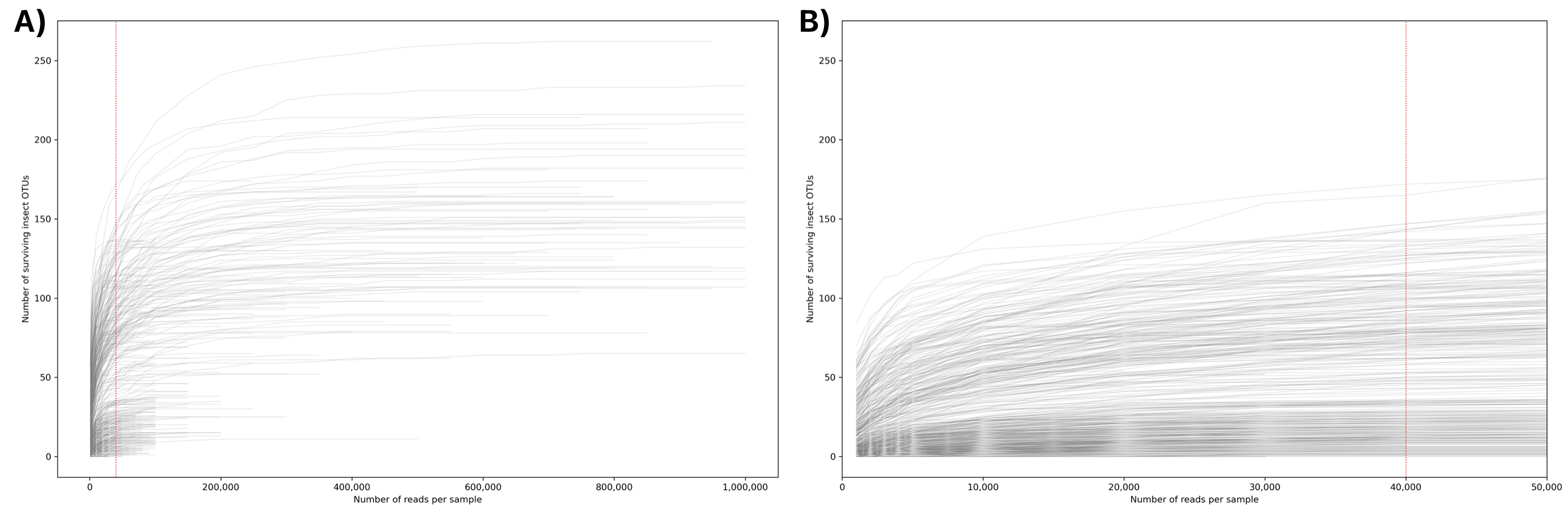

Figure S7. Rarefaction curves showing OTU richness as a function of sequencing depth for all samples. Panel A displays the full range of sequencing depths (up to 1,000,000 reads), whereas Panel B provides an enlarged view of the lower sequencing-depth range (up to 50,000 reads). The vertical red line in Panel B indicates the selected rarefaction threshold of 40,000 reads per sample, which was used to generate the rarefied OTU table for downstream diversity analyses.

Tables

Table S1. GPS of sampling locations.

| **Location** | **Longitude** | **Latitude** |
| --- | --- | --- |
| 1A | 17.90 | 60.11 |
| 1B | 17.87 | 60.18 |
| 1C | 17.82 | 60.04 |
| 2A | 17.81 | 59.51 |
| 2B | 17.63 | 59.63 |
| 2C | 17.71 | 59.55 |
| 3A | 13.00 | 59.34 |
| 3B | 13.22 | 59.46 |
| 3C | 13.01 | 59.46 |
| 4A | 15.07 | 58.95 |
| 4B | 15.15 | 58.94 |
| 4C | 15.25 | 58.86 |
| 5A | 16.72 | 59.15 |
| 5B | 16.55 | 59.25 |
| 5C | 16.49 | 59.20 |
| 6A | 14.13 | 58.56 |
| 6B | 14.30 | 58.59 |
| 6C | 14.30 | 58.53 |
| 7A | 12.35 | 58.10 |
| 7B | 12.41 | 58.07 |
| 7C | 12.64 | 58.06 |
| 8A | 12.98 | 57.42 |
| 8B | 13.02 | 57.43 |
| 8C | 12.81 | 57.40 |
| 9A | 14.33 | 57.69 |
| 9B | 14.31 | 57.55 |
| 9C | 14.06 | 57.62 |
| 10A | 16.02 | 58.17 |
| 10B | 15.90 | 58.22 |
| 10C | 15.89 | 58.15 |
| 11A | 15.68 | 57.52 |
| 11B | 15.65 | 57.37 |
| 11C | 15.51 | 57.41 |
| 12A | 13.77 | 57.07 |
| 12B | 13.85 | 57.13 |
| 12C | 13.88 | 57.02 |
| 13A | 13.43 | 56.51 |
| 13B | 13.60 | 56.55 |
| 13C | 13.61 | 56.47 |
| 14A | 14.87 | 56.91 |
| 14B | 14.94 | 56.87 |
| 14C | 14.96 | 56.98 |
| 15A | 15.81 | 56.33 |
| 15B | 15.72 | 56.28 |
| 15C | 15.75 | 56.33 |

Table S2. Dunn’s post-hoc pairwise comparisons for insect reads ratio (IRR, the percentage of insect reads to total reads) and insect OTUs ratio (IOR, OTU richness to total reads) with Benjamini-Hochberg corrected p-values. Significance levels are denoted as follows: ** (p ≤ 0.01), * (0.01 < p ≤ 0.05).

|  |  | **anthill** | **MT** | **sediment** | **soil** | **spiderweb** | **water** |
| --- | --- | --- | --- | --- | --- | --- | --- |
| **IRR** | | Kruskal-Wallis chi-squared = 235.3845, df = 6, p-value < 0.01 | | | | | |
| **MT** | Z-score | -6.6974 |  |  |  |  |  |
|  | P-value | 0.0000** |  |  |  |  |  |
| **sediment** | Z-score | -3.1104 | 5.9483 |  |  |  |  |
|  | P-value | 0.0015** | 0.0000** |  |  |  |  |
| **soil** | Z-score | -1.0374 | 12.5402 | 4.7079 |  |  |  |
|  | P-value | 0.1573 | 0.0000** | 0.0000** |  |  |  |
| **spiderweb** | Z-score | -2.8028 | 6.0494 | 0.3987 | -3.8133 |  |  |
|  | P-value | 0.0038** | 0.0000** | 0.3451 | 0.0001** |  |  |
| **water** | Z-score | -5.3365 | 2.2568 | -3.6915 | -9.5686 | -3.9054 |  |
|  | P-value | 0.0000** | 0.0258* | 0.0002** | 0.0000** | 0.0001** |  |
| **deadwood** | Z-score | -1.5908 | -8.4682 | -2.5200 | 1.3900 | -1.9951 | 6.2114 |
|  | P-value | 0.0651 | 0.0000** | 0.0082** | 0.0909 | 0.0284* | 0.0000** |
| **IOR** | | Kruskal-Wallis chi-squared = 226.3643, df = 6, p-value < 0.01 | | | | | |
| **MT** | Z-score | -4.3994 |  |  |  |  |  |
|  | P-value | 0.0001** |  |  |  |  |  |
| **sediment** | Z-score | -2.5820 | 3.0138 |  |  |  |  |
|  | P-value | 0.0074** | 0.0021** |  |  |  |  |
| **soil** | Z-score | 0.2784 | 10.1655 | 6.1972 |  |  |  |
|  | P-value | 0.4099 | 0.0000** | 0.0000** |  |  |  |
| **spiderweb** | Z-score | -1.3309 | 4.7931 | 1.9301 | -3.2320 |  |  |
|  | P-value | 0.1132 | 0.0000** | 0.0352* | 0.0012** |  |  |
| **water** | Z-score | -0.0704 | -1.1126 | -4.1264 | -11.6305 | -5.8501 |  |
|  | P-value | 0.0000** | 0.1469 | 0.0000** | 0.0000** | 0.0000** |  |
| **deadwood** | Z-score | -1.2152 | -5.2804 | -2.2667 | 3.2126 | -1.3309 | -6.3931 |
|  | P-value | 0.1308 | 0.0000** | 0.0164* | 0.0012** | 0.1132 | 0.0000** |

Table S3. Dunn’s post-hoc pairwise comparisons for insect Shannon and Simpson diversity indices with Benjamini-Hochberg corrected p-values. Significance levels are denoted as follows: ** (p ≤ 0.01), * (0.01 < p ≤ 0.05).

|  |  | | **anthill** | | **MT** | | **sediment** | | **soil** | **spiderweb** | | **water** | |
| --- | --- | --- | --- | --- | --- | --- | --- | --- | --- | --- | --- | --- | --- |
| **Shannon** | | | Kruskal-Wallis chi-squared = 160.08, df = 6, p-value = 0 | | | | | | | | | | |
| **MT** | | Z-score | -2.8833 |  | |  | |  | | |  | |  |
|  | | P-value | 0.0038** |  | |  | |  | | |  | |  |
| **sediment** | | Z-score | -2.4462 | 0.7476 | |  | |  | | |  | |  |
|  | | P-value | 0.0126* | 0.2513 | |  | |  | | |  | |  |
| **soil** | | Z-score | 0.2403 | 8.2509 | | 6.4081 | |  | | |  | |  |
|  | | P-value | 0.4253 | 0.0000** | | 0.0000** | |  | | |  | |  |
| **spiderweb** | | Z-score | -1.2394 | 3.1302 | | 2.2242 | | -3.5519 | | |  | |  |
|  | | P-value | 0.1329 | 0.0018** | | 0.0196* | | 0.0005** | | |  | |  |
| **water** | | Z-score | -3.6335 | -1.5554 | | -2.1744 | | -9.7265 | | | -4.4914 | |  |
|  | | P-value | 0.0004** | 0.0786 | | 0.0208* | | 0.0000** | | | 0.0000** | |  |
| **deadwood** | | Z-score | -1.2213 | -3.4264 | | -2.4313 | | 3.8445 | | | -0.0749 | | -4.8572 |
|  | | P-value | 0.1295 | 0.0007** | | 0.0122* | | 0.0002** | | | 0.4702 | | 0.0000** |
| **Simpson** | | | Kruskal-Wallis chi-squared = 147.1483, df = 6, p-value = 0 | | | | | | | | | | |
| **MT** | | Z-score | -2.6765 |  | |  | |  | | |  | |  |
|  | | P-value | 0.0071** |  | |  | |  | | |  | |  |
| **sediment** | | Z-score | -2.4917 | 0.2636 | |  | |  | | |  | |  |
|  | | P-value | 0.0095** | 0.4158 | |  | |  | | |  | |  |
| **soil** | | Z-score | 0.3081 | 7.8738 | | 6.6691 | |  | | |  | |  |
|  | | P-value | 0.4189 | 0.0000** | | 0.0000** | |  | | |  | |  |
| **spiderweb** | | Z-score | -1.0928 | 3.0189 | | 2.5768 | | -3.3487 | | |  | |  |
|  | | P-value | 0.1601 | 0.0027** | | 0.0081** | | 0.0012** | | |  | |  |
| **water** | | Z-score | -3.3003 | -1.2983 | | -1.4659 | | -9.0482 | | | -4.1421 | |  |
|  | | P-value | 0.0013** | 0.1274 | | 0.0999 | | 0.0000** | | | 0.0001** | |  |
| **deadwood** | | Z-score | -1.1640 | -3.1182 | | -2.6294 | | 3.8638 | | | 0.1021 | | -4.3031 |
|  | | P-value | 0.1510 | 0.0021** | | 0.0075** | | 0.0002** | | | 0.4593 | | 0.0000** |

Table S4. Dunn’s post-hoc pairwise comparisons for beta diversity dispersion with Benjamini-Hochberg corrected p-values. Significance levels are denoted as follows: ** (p < 0.01), * (0.01 < p < 0.05).

|  |  | | **anthill** | | **MT** | | **sediment** | | **soil** | | **spiderweb** | | | **water** |
| --- | --- | --- | --- | --- | --- | --- | --- | --- | --- | --- | --- | --- | --- | --- |
|  | | | | Kruskal-Wallis chi-squared = 188.24, df = 6, p-value = 0 | | | | | | | | | | |
| **MT** | | Z-score | | -1.7203 | |  | |  | |  | |  |  | |
|  | | P-value | | 0.0560 | |  | |  | |  | |  |  | |
| **sediment** | | Z-score | | -1.8971 | | -0.2932 | |  | |  | |  |  | |
|  | | P-value | | 0.0405* | | 0.3847 | |  | |  | |  |  | |
| **soil** | | Z-score | | -4.8959 | | -6.0770 | | -5.6908 | |  | |  |  | |
|  | | P-value | | 0.0000** | | 0.0000** | | 0.0000** | |  | |  |  | |
| **spiderweb** | | Z-score | | -3.9449 | | -3.6255 | | -3.3470 | | 0.9653 | |  |  | |
|  | | P-value | | 0.0001** | | 0.0003** | | 0.0007** | | 0.1848 | |  |  | |
| **water** | | Z-score | | 0.8285 | | 4.2268 | | 4.5200 | | 11.6424 | | 7.6408 |  | |
|  | | P-value | | 0.2139 | | 0.0000** | | 0.0000** | | 0.0000** | | 0.0000** |  | |
| **deadwood** | | Z-score | | -2.6392 | | 1.5237 | | 1.2305 | | -4.0706 | | -2.1780 | 5.7505 | |
|  | | P-value | | 0.0067** | | 0.0788 | | 0.1275 | | 0.0001** | | 0.0221* | 0.0000** | |

Table S5. Results of PERMANOVA testing the effects of substrate and location on insect order composition, and pairwise PERMANOVA between substrates with Benjamini-Hochberg corrected p-values, based on relative OTU richness. Significance levels are denoted as follows: ** (p ≤ 0.01), * (0.01 < p ≤ 0.05).

| **Factor** | | | **Df** | | **Sum of Squares** | | **R^2^** | | **F-Value** | | **P-value** | | **Adjusted p-value** | | | **Significance** | |
| --- | --- | --- | --- | --- | --- | --- | --- | --- | --- | --- | --- | --- | --- | --- | --- | --- | --- |
| PERMANOVA | | | | | | | | | | | | | | | | | |
| **Substrate** | | 6 | | 13.148 | | 0.1749 | | 13.7981 | | 0.001 | |  | | | ** | | |
| **Location** | | 44 | | 13.755 | | 0.18297 | | 1.9683 | | 0.001 | |  | | | ** | | |
| **Substrate: Location** | | 195 | | 20.796 | | 0.27663 | | 0.6715 | | 1 | |  | | |  | | |
| **Residual** | | 173 | | 27.476 | | 0.36549 | |  | |  | |  | | |  | | |
| **Total** |  | 418 | | 75.174 | | 1 | |  | |  | |  | | |  | | |
| Pairwise comparisons | | | | | | | | | | | | | | | | | |
| **substrate1** | **substrate2** |  | |  | |  | |  | | |  | | |  | | |  |
| **MT** | **sediment** | 1 | | 0.196 | | 6.913 | | 0.082 | | | 0.001 | | | 0.002 | | | ** |
| **MT** | **soil** | 1 | | 5.184 | | 26.534 | | 0.093 | | | 0.001 | | | 0.002 | | | ** |
| **MT** | **spiderweb** | 1 | | 1.828 | | 22.377 | | 0.223 | | | 0.001 | | | 0.002 | | | ** |
| **MT** | **water** | 1 | | 0.068 | | 6.238 | | 0.071 | | | 0.001 | | | 0.002 | | | ** |
| **MT** | **deadwood** | 1 | | 0.791 | | 15.568 | | 0.150 | | | 0.001 | | | 0.002 | | | ** |
| **MT** | **anthill** | 1 | | 0.034 | | 2.189 | | 0.043 | | | 0.086 | | | 0.095 | | |  |
| **sediment** | **soil** | 1 | | 3.216 | | 15.466 | | 0.059 | | | 0.001 | | | 0.002 | | | ** |
| **sediment** | **spiderweb** | 1 | | 1.420 | | 12.987 | | 0.162 | | | 0.001 | | | 0.002 | | | ** |
| **sediment** | **water** | 1 | | 0.189 | | 7.281 | | 0.093 | | | 0.001 | | | 0.002 | | | ** |
| **sediment** | **deadwood** | 1 | | 0.403 | | 5.723 | | 0.069 | | | 0.003 | | | 0.004 | | | ** |
| **sediment** | **anthill** | 1 | | 0.054 | | 1.187 | | 0.030 | | | 0.278 | | | 0.278 | | |  |
| **soil** | **spiderweb** | 1 | | 2.338 | | 10.436 | | 0.040 | | | 0.001 | | | 0.002 | | | ** |
| **soil** | **water** | 1 | | 4.718 | | 23.744 | | 0.086 | | | 0.001 | | | 0.002 | | | ** |
| **soil** | **deadwood** | 1 | | 3.143 | | 15.114 | | 0.055 | | | 0.001 | | | 0.002 | | | ** |
| **soil** | **anthill** | 1 | | 0.549 | | 2.407 | | 0.011 | | | 0.073 | | | 0.085 | | |  |
| **spiderweb** | **water** | 1 | | 1.577 | | 18.808 | | 0.207 | | | 0.001 | | | 0.002 | | | ** |
| **spiderweb** | **deadwood** | 1 | | 0.671 | | 5.440 | | 0.065 | | | 0.003 | | | 0.004 | | | ** |
| **spiderweb** | **anthill** | 1 | | 0.371 | | 2.444 | | 0.059 | | | 0.057 | | | 0.070 | | |  |
| **water** | **deadwood** | 1 | | 0.692 | | 13.708 | | 0.143 | | | 0.001 | | | 0.002 | | | ** |
| **water** | **anthill** | 1 | | 0.053 | | 5.249 | | 0.109 | | | 0.005 | | | 0.007 | | | ** |
| **deadwood** | **anthill** | 1 | | 0.140 | | 1.703 | | 0.034 | | | 0.174 | | | 0.183 | | |  |

Table S6. Dunn’s post-hoc pairwise comparisons for order composition dispersion with Benjamini-Hochberg corrected p-values, based on relative OTU richness data. Significance levels are denoted as follows: ** (p ≤ 0.01), * (0.01 < p ≤ 0.05).

|  |  | | **anthill** | **MT** | **sediment** | **soil** | **spiderweb** | **water** |
| --- | --- | --- | --- | --- | --- | --- | --- | --- |
|  | | | Kruskal-Wallis chi-squared = 276.8758, df = 6, p-value = 0 | | | | | |
| **MT** | | Z-score | 0.5941 |  |  |  |  |  |
|  | | P-value | 0.2900 |  |  |  |  |  |
| **sediment** | | Z-score | -0.7726 | -2.6420 |  |  |  |  |
|  | | P-value | 0.2565 | 0.0067** |  |  |  |  |
| **soil** | | Z-score | -4.0864 | -11.8926 | -7.3105 |  |  |  |
|  | | P-value | 0.0001** | 0.0000** | 0.0000** |  |  |  |
| **spiderweb** | | Z-score | -2.7747 | -6.5855 | -3.6706 | 2.5531 |  |  |
|  | | P-value | 0.0053** | 0.0000** | 0.0000** | 0.0080** |  |  |
| **water** | | Z-score | 10.6485 | 0.1196 | 2.6701 | 11.3517 | 6.4869 |  |
|  | | P-value | 0.2855 | 0.4524 | 0.0066** | 0.0000** | 0.0000** |  |
| **deadwood** | | Z-score | -1.8342 | 5.0060 | 2.0025 | -5.4548 | -1.9029 | 4.9435 |
|  | | P-value | 0.0412* | 0.0000** | 0.0317* | 0.0000** | 0.0374* | 0.0000** |

Table S7. Results of PERMANOVA testing the effects of substrate and location on insect order composition, and pairwise PERMANOVA between substrates with Benjamini-Hochberg corrected p-values, based on relative insect read abundance data. Significance levels are denoted as follows: ** (p ≤ 0.01), * (0.01 < p ≤ 0.05).

| **Factor** | | | **Df** | | **Sum of Squares** | | **R^2^** | | **F-Value** | **P-value** | **Adjusted p-value** | | **Significance** | |
| --- | --- | --- | --- | --- | --- | --- | --- | --- | --- | --- | --- | --- | --- | --- |
| **PERMANOVA** | | | | | | | | | | | | | | |
| Substrate | | 6 | | 35.363 | | 0.213 | | 20.289 | | 0.001 | |  | | ** |
| Location | | 44 | | 25.107 | | 0.151 | | 1.964 | | 0.001 | |  | | ** |
| Substrate: Location | | 195 | | 55.135 | | 0.332 | | 0.973 | | 0.785 | |  | |  |
| Residual | | 173 | | 50.257 | | 0.303 | |  | |  | |  | |  |
| Total |  | 418 | | 165.862 | | 1 | |  | |  | |  | |  |
| **Pairwise comparisons** | | | | | | | | | | | | | | |
| substrate1 | substrate2 |  | |  | |  | |  | |  | |  | |  |
| MT | sediment | 1 | | 7.366 | | 0.322 | | 36.495 | | 0.001 | | 0.001 | | ** |
| MT | soil | 1 | | 13.725 | | 0.128 | | 37.757 | | 0.001 | | 0.001 | | ** |
| MT | spiderweb | 1 | | 9.055 | | 0.331 | | 38.675 | | 0.001 | | 0.001 | | ** |
| MT | water | 1 | | 7.064 | | 0.383 | | 51.008 | | 0.001 | | 0.001 | | ** |
| MT | deadwood | 1 | | 14.020 | | 0.472 | | 78.809 | | 0.001 | | 0.001 | | ** |
| MT | anthill | 1 | | 4.235 | | 0.398 | | 32.400 | | 0.001 | | 0.001 | | ** |
| sediment | soil | 1 | | 3.446 | | 0.034 | | 8.763 | | 0.001 | | 0.001 | | ** |
| sediment | spiderweb | 1 | | 1.665 | | 0.072 | | 5.164 | | 0.002 | | 0.002 | | ** |
| sediment | water | 1 | | 1.865 | | 0.113 | | 9.009 | | 0.001 | | 0.001 | | ** |
| sediment | deadwood | 1 | | 3.012 | | 0.137 | | 12.212 | | 0.001 | | 0.001 | | ** |
| sediment | anthill | 1 | | 1.516 | | 0.135 | | 5.914 | | 0.004 | | 0.004 | | ** |
| soil | spiderweb | 1 | | 1.995 | | 0.020 | | 4.955 | | 0.001 | | 0.001 | | ** |
| soil | water | 1 | | 10.135 | | 0.098 | | 27.479 | | 0.001 | | 0.001 | | ** |
| soil | deadwood | 1 | | 5.323 | | 0.052 | | 14.124 | | 0.001 | | 0.001 | | ** |
| soil | anthill | 1 | | 1.610 | | 0.018 | | 4.007 | | 0.001 | | 0.001 | | ** |
| spiderweb | water | 1 | | 5.865 | | 0.252 | | 24.246 | | 0.001 | | 0.001 | | ** |
| spiderweb | deadwood | 1 | | 1.531 | | 0.066 | | 5.499 | | 0.001 | | 0.001 | | ** |
| spiderweb | anthill | 1 | | 1.072 | | 0.079 | | 3.353 | | 0.003 | | 0.003 | | ** |
| water | deadwood | 1 | | 10.362 | | 0.412 | | 57.376 | | 0.001 | | 0.001 | | ** |
| water | anthill | 1 | | 3.738 | | 0.402 | | 28.914 | | 0.001 | | 0.001 | | ** |
| deadwood | anthill | 1 | | 0.230 | | 0.023 | | 1.145 | | 0.270 | | 0.270 | |  |

Table S8. Mean of relative read abundance of different insect orders across substrates (%).

| **Order** | **MT** | **water** | **sediment** | **spiderweb** | **deadwood** | **anthill** | **soil** |
| --- | --- | --- | --- | --- | --- | --- | --- |
| **Coleoptera** | 0.00 | 0.10 | 0.12 | 1.93 | 1.93 | 0.14 | 15.12 |
| **Dermaptera** | 0.01 | 0.00 | 0.01 | 0.00 | 0.00 | 0.00 | 0.00 |
| **Diptera** | 94.35 | 95.47 | 92.23 | 63.07 | 90.43 | 97.26 | 51.20 |
| **Embioptera** | 0.00 | 0.00 | 1.07 | 0.00 | 0.00 | 0.00 | 0.00 |
| **Ephemeroptera** | 0.00 | 0.08 | 1.75 | 0.00 | 0.00 | 0.00 | 0.00 |
| **Hemiptera** | 1.15 | 0.98 | 3.58 | 5.50 | 2.53 | 1.33 | 27.44 |
| **Hymenoptera** | 0.07 | 0.02 | 0.02 | 1.47 | 0.09 | 0.00 | 0.79 |
| **Lepidoptera** | 0.06 | 0.03 | 0.00 | 4.83 | 0.00 | 0.00 | 0.12 |
| **Neuroptera** | 0.02 | 0.01 | 0.00 | 0.00 | 0.00 | 0.00 | 0.00 |
| **Odonata** | 0.00 | 0.01 | 0.00 | 0.00 | 0.00 | 0.00 | 0.00 |
| **Plecoptera** | 0.00 | 0.26 | 0.39 | 0.00 | 0.00 | 0.00 | 0.05 |
| **Psocodea** | 4.21 | 2.57 | 0.69 | 11.77 | 2.67 | 1.28 | 0.93 |
| **Raphidioptera** | 0.00 | 0.00 | 0.00 | 0.00 | 0.18 | 0.00 | 0.00 |
| **Thysanoptera** | 0.00 | 0.00 | 0.00 | 0.00 | 0.00 | 0.00 | 1.04 |
| **Trichoptera** | 0.11 | 0.48 | 0.12 | 10.37 | 0.39 | 0.00 | 2.77 |
| **Zoraptera** | 0.00 | 0.00 | 0.02 | 1.05 | 1.78 | 0.00 | 0.54 |

Table S9. Dunn’s post-hoc pairwise comparisons of assigned OTU proportion with Benjamini-Hochberg corrected p-values. Significance levels are denoted as follows: ** (p ≤ 0.01), * (0.01 < p ≤ 0.05).

|  |  | | **anthill** | | **MT** | | **sediment** | | **soil** | **spiderweb** | **water** | |
| --- | --- | --- | --- | --- | --- | --- | --- | --- | --- | --- | --- | --- |
| **family** | | | Kruskal-Wallis chi-squared = 34.2088, df = 6, p-value = 0 | | | | | | | | | |
| **MT** | | Z-score | -0.1680 |  | |  | |  | |  | |  |
|  | | P-value | 0.4333 |  | |  | |  | |  | |  |
| **sediment** | | Z-score | 1.3336 | 2.9201 | |  | |  | |  | |  |
|  | | P-value | 0.1368 | 0.0073** | |  | |  | |  | |  |
| **soil** | | Z-score | 0.7430 | 2.3214 | | -1.5333 | |  | |  | |  |
|  | | P-value | 0.2669 | 0.0266* | | 0.1096 | |  | |  | |  |
| **spiderweb** | | Z-score | 1.7106 | 3.6776 | | 0.6865 | | 2.4595 | |  | |  |
|  | | P-value | 0.0832 | 0.0006** | | 0.2721 | | 0.0209* | |  | |  |
| **water** | | Z-score | 0.2459 | 0.8266 | | -2.0573 | | -1.1475 | | -2.7831 | |  |
|  | | P-value | 0.4230 | 0.2681 | | 0.0416* | | 0.1758 | | 0.0094** | |  |
| **deadwood** | | Z-score | 2.1426 | -4.7634 | | -1.4993 | | -3.8044 | | -0.7782 | | -3.7636 |
|  | | P-value | 0.0375* | 0.0000** | | 0.1081 | | 0.0007** | | 0.2696 | | 0.0006** |
| **genus** | | | Kruskal-Wallis chi-squared = 50.3044, df = 6, p-value = 0 | | | | | | | | | |
| **MT** | | Z-score | -1.0209 |  | |  | |  | |  | |  |
|  | | P-value | 0.2482 |  | |  | |  | |  | |  |
| **sediment** | | Z-score | 0.5219 | 2.9695 | |  | |  | |  | |  |
|  | | P-value | 0.3326 | 0.0052** | |  | |  | |  | |  |
| **soil** | | Z-score | 0.9919 | 5.2110 | | 0.9725 | |  | |  | |  |
|  | | P-value | 0.2409 | 0.0000** | | 0.2316 | |  | |  | |  |
| **spiderweb** | | Z-score | 0.6185 | 3.1812 | | 0.1753 | | -0.7532 | |  | |  |
|  | | P-value | 0.3128 | 0.0031** | | 0.4304 | | 0.2788 | |  | |  |
| **water** | | Z-score | -0.9222 | 0.1794 | | -2.7085 | | -4.6826 | | -2.9107 | |  |
|  | | P-value | 0.2339 | 0.4503 | | 0.0089** | | 0.0000** | | 0.0054** | |  |
| **deadwood** | | Z-score | 1.4234 | -5.0389 | | -1.7054 | | -1.2691 | | -1.5322 | | -4.6762 |
|  | | P-value | 0.1476 | 0.0000** | | 0.1028 | | 0.1789 | | 0.1317 | | 0.0000** |
| **species** | | | Kruskal-Wallis chi-squared = 18.2974, df = 6, p-value = 0 | | | | | | | | | |
| **MT** | | Z-score | 0.2751 |  | |  | |  | |  | |  |
|  | | P-value | 0.4329 |  | |  | |  | |  | |  |
| **sediment** | | Z-score | 1.0266 | 1.4745 | |  | |  | |  | |  |
|  | | P-value | 0.2665 | 0.1474 | |  | |  | |  | |  |
| **soil** | | Z-score | 1.0064 | 1.8117 | | -0.2062 | |  | |  | |  |
|  | | P-value | 0.2538 | 0.1226 | | 0.4392 | |  | |  | |  |
| **spiderweb** | | Z-score | 0.7377 | 0.9158 | | -0.5342 | | -0.4970 | |  | |  |
|  | | P-value | 0.3023 | 0.2518 | | 0.3664 | | 0.3612 | |  | |  |
| **water** | | Z-score | -0.1854 | -0.9180 | | -2.2839 | | -2.8604 | | -1.7491 | |  |
|  | | P-value | 0.4265 | 0.2690 | | 0.0587 | | 0.0148* | | 0.1054 | |  |
| **deadwood** | | Z-score | 1.6887 | -2.9143 | | -1.2294 | | -1.9362 | | -1.8103 | | -3.7263 |
|  | | P-value | 0.1065 | 0.0187* | | 0.2090 | | 0.1110 | | 0.1054 | | 0.0020** |

Table S10. Unique insect species count exclusively found by substrates in different orders.

| **substrate** | **order** | **unique species** |
| --- | --- | --- |
| **MT** | Coleoptera | 8 |
|  | Diptera | 94 |
|  | Hemiptera | 11 |
|  | Hymenoptera | 31 |
|  | Lepidoptera | 4 |
|  | Neuroptera | 2 |
| **sediment** | Coleoptera | 1 |
|  | Diptera | 1 |
| **soil** | Coleoptera | 5 |
|  | Diptera | 5 |
|  | Hymenoptera | 1 |
| **spiderweb** | Coleoptera | 4 |
|  | Hemiptera | 1 |
|  | Lepidoptera | 1 |
|  | Psocodea | 1 |
| **water** | Coleoptera | 5 |
|  | Diptera | 15 |
|  | Ephemeroptera | 1 |
|  | Hemiptera | 1 |
|  | Lepidoptera | 4 |
|  | Odonata | 2 |
|  | Plecoptera | 1 |
|  | Trichoptera | 4 |
| **deadwood** | Coleoptera | 4 |
|  | Raphidioptera | 1 |

Supplementary Data (separate file)

Data S1. Taxonomic assignments of the 466 insect species recovered in this study and their presence–absence across substrate types.

Data S2. List of substrate-specific species and their taxonomic assignments.
